# Arousal elicits a brain-wide hemodynamic wave independent of locus coeruleus noradrenergic tone

**DOI:** 10.64898/2026.03.06.710089

**Authors:** Jose Maria Martinez de Paz, Johanna Luise Mayer, Paulina Wanken, Beatriz Rodrigues Apgaua, Aliya Ablitip, Leafy Behera, Émilie Macé

## Abstract

Arousal fluctuations during wakefulness have a major impact on physiology and behavior, including perception and task performance. Arousal is also known to be a strong modulator of neural activity, but the brain-wide spatiotemporal structure of this modulation is not fully characterized. We used functional ultrasound imaging to record brain-wide hemodynamics - a proxy for neural activity - in head-fixed mice during spontaneous and sensory-evoked arousal fluctuations, tracked via pupil diameter. Both conditions recruited a common brain-wide hemodynamic wave that followed a subcortex-to-cortex gradient. We then tested whether noradrenaline, widely associated with arousal, was necessary or sufficient to drive this wave. Sustained bidirectional optogenetic manipulations of locus coeruleus activity affected brain wide vascular signal amplitude but, surprisingly, left arousal-linked dynamics largely intact. Together, these results identify a common spatiotemporal motif of arousal that appears independent of noradrenergic tone.

## Introduction

Arousal is hypothesized to be a core, evolutionarily conserved brain function that enables animals to modulate their responsiveness to the external world according to their internal needs ^1^. As such, it can be defined as the degree of sensitivity to external stimuli, ranging from sleep and inattention on one side, to hypervigilance and hyperactivity on the other ^2,3^. We focus here on the middle range of that spectrum: moderate arousal fluctuations within wakefulness. These fluctuations have a profound impact on cognition and behavior, including task performance, perceptual abilities and muscle tone^4-6^. In parallel, arousal dynamics have a major effect on brain activity, such as strong changes in cortical synchronization^7-10^. In humans, arousal fluctuations strongly modulate global brain hemodynamics ^11,12.^ In rodents, large-scale neural recordings have also revealed that changes in arousal - tracked using behavioral proxies - explain large fractions of brain activity both at rest ^13-15^ and when the animal is engaged in a task ^16-19^.

Despite the widespread impact of arousal on brain activity, it remains unclear how these effects distribute across brain regions. Existing studies suggest regionally heterogeneous effects of arousal, but their results are technically constrained either to superficial cortex, in the case of widefield optical imaging ^15-16-18^ or to restricted sets of deep structures, for multielectrode recordings^20^. While whole-brain approaches such as functional magnetic resonance imaging (fl’v1RI) could in principle overcome this limitation, they typically require immobilization and are therefore biased towards sedated or low-arousal states ^21-23^. Consequently, and despite recent efforts to expand spatial coverage in awake mice ^20,24,^ a comprehensive characterization of brain-wide arousal dynamics during wakefulness is still lacking.

These limitations apply not only to spontaneous arousal fluctuations, but also to arousal responses elicited by external events. Arousal reliably increases in response to salient or surprising stimuli, as shown by a large body of work using air puffs, sounds, or oddball paradigms ^10,25-28^. Although the term “arousal” is used in both contexts, it is unclear whether spontaneous and evoked arousal events engage the same large-scale brain pathways. Recent studies suggest that evoked arousal activates cortical areas also involved in spontaneous arousal ^29^, but direct whole-brain comparisons between the two conditions are missing. Notably, such a comparison would offer the opportunity to dissect arousal mechanisms from two distinct experimental angles in order to isolate the common latent factors.

At the mechanistic level, arousal states rely on multiple neuromodulatory systems, including noradrenaline^2-30^ acetylcholine^31-32,^ serotonin ^33,34,^ hypocretin/orexin ^35^, etc. Among those, the locus coeruleus-noradrenaline (LC-NA) system has been particularly well studied ^3,35,37^. Noradrenergic tone closely tracks behavioral state - as reflected by pupil size^38-40^ - and increases in response to salient external stimuli^41-43^ thus positioning the LC-NA system as a plausible mediator of both spontaneous and evoked arousal. Experimental manipulations of noradrenergic tone have also been shown to alter pupil size^41,44,45^ and broadly modulate brain activity^46-49^. However, whether noradrenergic tone causally contributes to the brain-wide effects of arousal remains largely unresolved.

The present study addresses these gaps by providing a whole-brain characterization of both spontaneous and evoked arousal under varying LC activity tone. We recorded brain-wide activity from awake, head-fixed mice using volumetric functional ultrasound imaging (fUSi) ^50,51^. Functional ultrasound imaging is a neuroimaging method that measures blood volume changes, which correlate with low-frequency neuronal activity^52^, and enables a spatial resolution of ∼300 µm and a temporal resolution of 2.5 Hz for brain-wide recordings ^50^ We observed a robust and recurring spatiotemporal hemodynamic wave that emerged both during spontaneous pupil dilations and in response to arousing stimuli (mild air puffs or white noise). This wave was recoverable via data-driven decomposition of resting-state activity and predictive of evoked arousal responses, consistent with a shared latent arousal structure. Finally, bidirectional manipulation of tonic noradrenergic levels via sustained optogenetic stimulation of the locus coeruleus broadly affected fUS signal amplitude, but did not alter the structure of the arousal wave during either spontaneous or evoked transitions. Together, these findings provide new insights into the brain-wide effects of arousal by identifying a large-scale arousal-related hemodynamic wave that persists under tonic modulation of noradrenergic neurons, suggesting that other neuromodulators may play a more prominent role in driving arousal dynamics during wakefulness.

## Results

### Spontaneous arousal fluctuations in awake mice heterogeneously affect whole-brain hemodynamics

To investigate the relationship between spontaneous arousal fluctuations and whole-brain activity, we performed volumetric fUSi on head-fixed mice within a virtual burrow assay (VBA)^53^ while simultaneously monitoring pupil size, orofacial movements and the burrow speed in the absence of stimuli **(Fig. la, Methods)**. The VBA offers the mice a shelter to rest while simultaneously allowing them to move back and forth during higher arousal states. Brain data collection occurred through a COMBO cranial window^54^ that ranged from the caudal end of the olfactory bulb to the rostral edge of the cerebellum **(Fig. 1b)**. The behavioral proxies of arousal spontaneously fluctuated in strong synchrony with each other, with face and burrow motion happening on average 0.86 ± 0.32 s before maximum pupil dilation **(Fig. lc,d** and **Supplementary Fig. la, b)**. Because pupil fluctuations are well-documented to reflect internal arousal dynamics ^55^, we focused on pupil size as our main proxy for arousal fluctuations. Cross-correlation analysis between pupil size and voxel-wise fUSi time courses revealed a robust, spatially heterogeneous brain-wide pattern of positive correlations whose strength varied across brain voxels **(Fig. le)**. These correlations were strongest in cortex, thalamus and midbrain but substantial variability existed at finer spatial scales within these structures, indicating high heterogeneity both across and within anatomical units **(Fig. lf)**.

**Figure 1.**
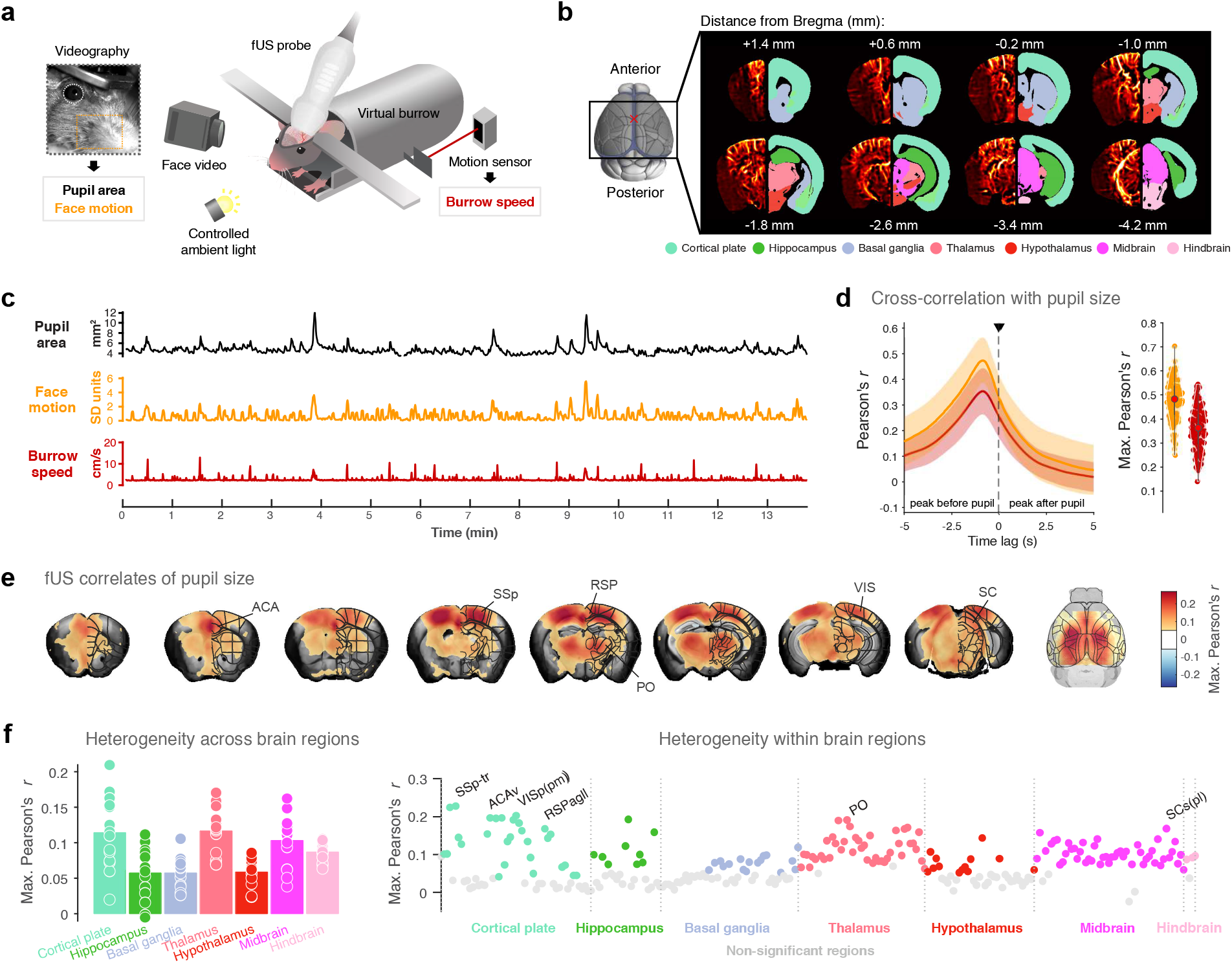
Spontaneous arousal fluctuations span heterogeneous brain-wide correlates. **a.** Schematic of the experimental setup, combining head-fixation inside a moving virtual burrow, functional ultrasound imaging (fUSi), videographic tracking of face and pupil, and a motion sensor for burrow position. The setup was placed inside a sound-proof box which in turn was over an anti-vibration table. **b**. Imaging field-of-view along the rostrocaudal axis and coronal slices of a representative power Doppler volumetric image with overlaid anatomical brain regions, as defined in the Allen Brain Atlas Common Coordinate Framework. Bregma is marked with a red X. The most lateral regions, such as temporal, insular and piriform cortices, lay largely outside the imaging window. **c**. Example time series of behavioral metrics during a representative stimulus-free recording, showing spontaneous, aligned fluctuations. **d**. Left: average cross-correlation between burrow or face motion and the pupil trace (*n* = 105 behavioral traces). Right: peak correlation distributions across sessions. Line plot represents mean cross-correlation; shaded area represents ± standard deviation. **e**. Left: brain-wide maximum correlation map of pupil size with fUS signal, with brain regions overlaid (FDR < 0.15, *n* = 105 sessions, *N* = 15 mice). Shade over the background image represents window coverage. Right: cortical view of the same map. **f**. Left: average maximum correlation values across the major anatomical brain areas shown in **b**. Right: average maximum correlation values across the fine brain regions shown in the overlay from **e**. Significant regions are defined as those with >55% significant voxels in e. ACA: anterior cingulate area; SSP: primary somatosensory cortex; RSP: retrosplenial cortex; PO: pulvinar nucleus; VIS: primary visual cortex; SC: superior colliculus.

Repeating the analysis using facial and burrow motion instead of pupil size yielded almost identical maps **(Supplementary Fig. lc,d)**, raising the possibility that the observed brain-wide pattern could reflect motor activity. To disentangle the unique contributions of each behavioral proxy, we employed partial correlation analysis. While partial correlates of all three behavioral traces remained present in the cortex, suggesting additive effects, volumetric fUS revealed that only the partial correlates of pupil size extended into the thalamus and midbrain, closely matching the full correlation maps **(Supplementary Fig. le,f)**, thus indicating that the brain-wide pattern cannot be explained by motion alone. Additionally, to exclude a confounding effect of arousal-independent pupil changes, such as the light reflex, we exposed mice to LED-driven light ramps inducing slow cycles of pupil constriction and dilation **(Supplementary Fig. 2a,b)**. This manipulation failed to evoke significant correlations outside the prelimbic cortex **(Supplementary Fig. 2c)**, arguing against a purely mechanistical origin of the spontaneous pupil-brain correlations. Together, these results capture a robust, brain-wide hemodynamic pattern associated with spontaneous arousal fluctuations during wakefulness that cannot be accounted for solely by concurrent movement or light-driven pupil dynamics.

### Spontaneous and evoked arousal events engage similar spatiotemporal dynamics of activation

Since arousal is known to increase in response to sensory stimuli ^10,27^, we sought to identify context-independent arousal effects by assessing whether spontaneous and evoked arousal transitions engage the same brain structures. To this end, we designed an experimental paradigm to compare spontaneous and evoked arousal at the whole-brain level. Mice were recorded in the virtual burrow while brief air puffs (0.2 s) were delivered to their left cheek **(Fig. 2a)**. These air puffs were calibrated to be arousing but mild, in order to minimize burrow motion and associated motor confounders. Each imaging session consisted of a stimulus-free baseline followed by a period of pseudorandom air-puff delivery (19.9 per session on average), enabling paired analysis between spontaneous and evoked arousal conditions **(Fig. 2b)**. Spontaneous arousal events were defined as sudden, isolated and large pupil dilations during the baseline period (see **Methods)**. These spontaneous pupil events typically occurred every 80 s together with burrow motion, whereas air puffs reliably evoked pupil dilation with only minimal accompanying movement and no habituation across trials **(Supplementary Fig**. 3).

**Figure 2.**
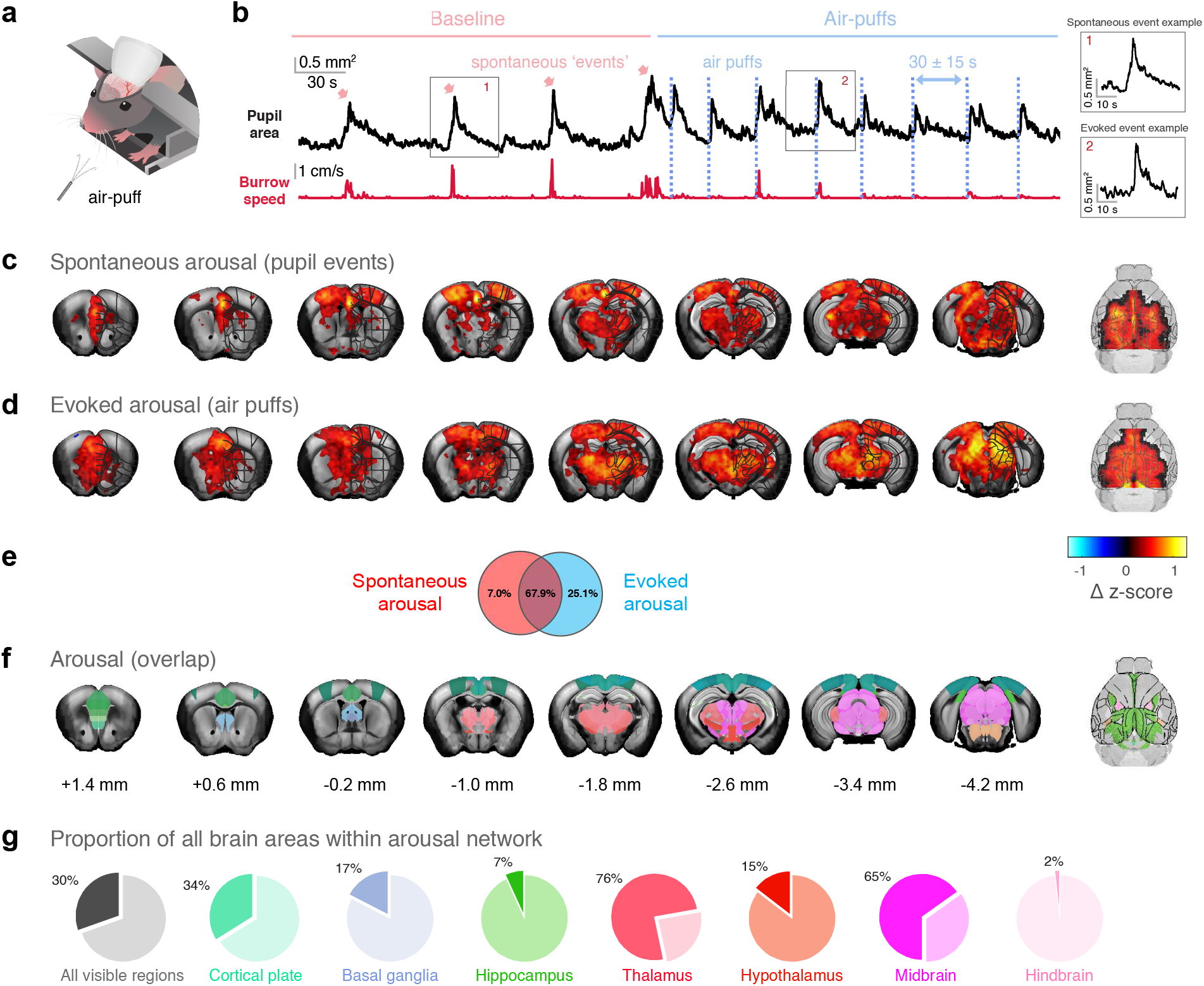
Arousal regions activated by both spontaneous and evoked arousal events. **a.** Schematic of the air puff delivery to evoke sensory-driven arousal. **b**. Left: example pupil and burrow speed recording from one imaging session, showing the spontaneous events and evoked air puff responses. Right: detail of the pupil trace upon spontaneous and evoked arousal events. c. Left: average brain-wide fUS response amplitude for spontaneous pupil events (FDR<0.15, *n* = 410 events, *N* = 7 mice), overlaid with sub-regions delimitations. The fUS response amplitude is calculated as the maximum absolute value after the timepoint of maximum pupil change-rate. Right: cortical view of the same maps. **d**. As **c** but for air puffs (FDR<0.15, *n* = 419 puffs, 7 *N* = mice). The fUS response amplitude is calculated from the time of the air puff delivery. e. Venn diagram showing the percentage of significant brain regions from either spontaneous or evoked maps that appear in both conditions. Significant regions are defined as those with >55% significant voxels in d. A full list of the shared arousal regions is presented in the Supplementary Table 1. **f**. Left: whole-brain sub regions that are significantly active in response to both spontaneous pupil events and air puffs (same ones as in e). Right: cortical view of the same map. **g**. Pie charts showing which percentage of the total brain regions respond to both spontaneous and evoked arousal.

For both spontaneous and evoked arousal conditions, we extracted voxel-wise fUS signals time-locked to arousal events **(Methods)**. This analysis revealed a structured, large-scale “wave-like” propagation of fUS activity from midbrain to frontal cortex in both conditions **(Supplementary Movie 1, 2)**. Fiber photometry recordings in the caudal cingulate cortex, one of the common active areas in both waves, showed calcium activity increases across conditions, thus discarding a purely hemodynamic explanation of these responses **(Supplementary Fig. 4)**. Static peak-response maps derived from the waves (FDR<0.15) showed a similar spatial organization for both spontaneous and evoked arousal events (spatial Pearson’s *r* = 0.56; **Fig. 2c,d)**, and closely resembled the pupil correlates from **(Fig. le)**. Notably, evoked arousal responses exhibited stronger activation in thalamic and midbrain regions compared to spontaneous pupil events. Both the temporal evolution of the fUS signal and the peak-response maps were reproducible across animals, with evoked arousal responses showing greater inter-individual consistency **(Supplementary Fig. 5)**. We confirmed the generalizability of the evoked arousal response with a different sensory modality, namely brief bouts of white noise, which also elicited a fUS pattern similar to that evoked by air puffs (spatial Pearson’s *r* = 0.67; **Supplementary Fig. 6a)**. Moreover, to determine whether the enhanced thalamic and midbrain responses after air puffs reflected sensory processing rather than arousal per se, we collected a subset of air puff trials under anesthesia. In this condition, both brief (0.2 s) and sustained (10 s) air puffs evoked fUS responses restricted to thalamus and superior colliculus, plus the contralateral barrel cortex in the case of sustained stimuli **(Supplementary Movie 3, Supplementary Fig. 6b-c)**. These maps indicate that thalamus and superior colliculus capture a sensory component of the air puff response, whereas the broader brain-wide response likely reflects arousal-related activity beyond sensory processing.

Upon parcellating the static maps, we found that a large fraction (67.9%) of brain regions significantly active during either spontaneous or evoked arousal overlapped across both conditions **(Fig. 2e)**. These core arousal regions (depicted in **Fig. 2f** and listed in **Supplementary table 1)** comprised most of the thalamus as well as large portions of the mid brain and cortex **(Fig. 2g)**. Together, these results support the existence of a specific arousal response independent of the source of arousal (i.e. internally- or externally-driven) that cannot be explained by motor or sensory responses alone.

Having established a shared spatial organization, we next sought to test whether the temporal dynamics of these large-scale activations were consistent across both conditions. To this end, we extracted the voxel-wise time till maximum absolute activation around arousal events to generate delay maps for spontaneous and evoked conditions **(Fig. 3a, Methods)**. While spontaneous arousal spanned over a longer timeframe than evoked arousal (5.70 vs. 4.37 seconds, respectively), the spatial organization of the delay maps was remarkably conserved, with cortical regions consistently peaking later than subcortical regions in both conditions. To facilitate comparisons, we grouped the voxel-wise delays into anatomical regions belonging to the core arousal network **(Fig. 3b**). The region-wise relative delays were highly correlated across spontaneous and evoked conditions **(Fig. 3c)**, confirming a common activation sequence that originated in caudal and ventral regions, namely the midbrain, and propagated frontally and laterally along the cortex. Since the temporal profile of subcortical regions activity differed across conditions, with evoked responses being temporally sharper than spontaneous ones **(Fig. 3d)**, we confirmed the relative temporal ordering with a timescale-independent cross-correlation analysis **(Methods)** that yielded nearly identical subcortex-to-cortex activation sequences in both conditions **(Fig. 3e,f)**. Notably, while spontaneous and evoked arousal differ in temporal profiles, the shared brain-wide sequence of regional activation strongly indicated they reflect a common underlying dynamic.

**Figure 3.**
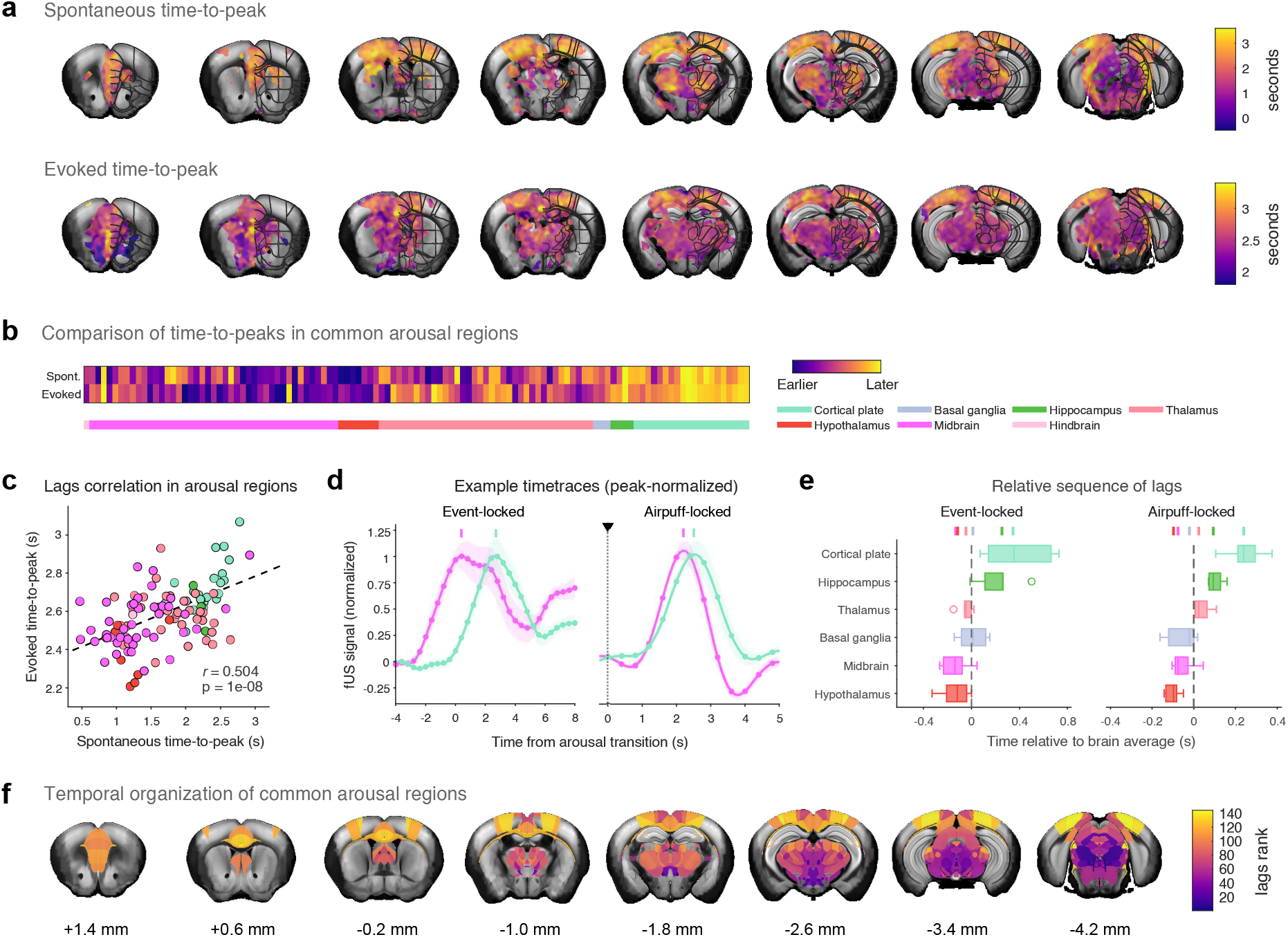
Brain regions modulated by arousal follow a consistent brainstem-to-cortex temporal gradient of activation. **a.** Coronal slices of the average brain-wide fUS response time-to-peak for spontaneous pupil events *(top)* and for air puffs *(bottom)*. **b**. Heatmap of each region time-to-peak for spontaneous and evoked arousal. Regions are defined according to the Allen Brain Atlas Common Coordinate Framework. **c**. Scatterplot comparing each brain region time-to-peak for spontaneous vs. evoked arousal. Black line indicates the best linear fit. **d**. Average time courses of representative brain macro-regions for spontaneous (*left)* and evoked (*right)* arousal events, with peaks marked as solid lines above. Macro-regions are defined as the broadest anatomical categories from the Allen Brain Atlas Common Coordinate Framework. **e**. Relative sequence of activation of each brain macro-regions for spontaneous (*left)* and evoked (*right)* arousal, calculated as the time-to-maximum correlation of each macro-region with the global mean, with the median lags marked as solid lines above. Black dashed line represents the global mean and defines zero. **f**. Coronal slices showing the average relative (ranked) sequence of activation across brain regions.

Finally, to assess whether the observed timing differences reflected neural population dynamics rather than region-specific hemodynamic coupling, we leveraged an independent fiber photometry dataset acquired in awake mice within the VBA. Fiber photometry recordings in regions from the fUSi-defined arousal network showed calcium activity increases around pupil events, with region-specific fiber photometry delays between calcium signal and pupil peak dilation closely matching the fUSi-derived times until maximum activation **(Supplementary Fig. 7)**. Together, these results provide independent validation of the conserved temporal gradient identified with fUSi and support the interpretation that the observed propagation reflects genuine arousal-related neural dynamics rather than vascular confounds.

### Spontaneous and evoked arousal transitions share low-dimensional subspace

Low-dimensional representations of brain activity in selected regions often reflect behavioral proxies of arousal ^19,56,57.^ Given the strong effects of arousal events on fUS signal, we asked whether this influence was general and salient enough to be captured by dimensionality reduction of spontaneous whole-brain data. For this, we performed group principal component analysis (PCA) on independent stimulus-free (i.e. “resting state”) fUSi recordings **(Fig. 4a, Methods)**. In this training dataset, the first two principal components (PCs) captured the majority of resting-state group variance (63.4%), with next components only providing minimal gains **(Fig. 4b)**. These first two PCs showed strong spatial overlap with the core arousal regions identified in our voxel-wise analysis **(Fig. 2f):** the first was a cortex-dominated component (PCl^cort^) and the second, a thalamus- and midbrain-dominated component (PC2^subcort^) **(Fig. 4c)**. Back-projecting these spatial components onto session-specific fUS data to get their temporal weights **(Methods)** revealed that both PCs were highly correlated with pupil size fluctuations. Moreover, PC2^subcort^ temporally preceded PCl ^cort^ relative to the pupil cycle **(Fig. 4d)**, matching the subcortex-to-cortex progression identified in our time-locked analyses **(Fig. 3)**. Together, these results indicate that low-dimensional arousal-related activity accounts for the majority of fUS recordings variance during resting state, to the point that it can be robustly recovered from spontaneous whole-brain fluctuations. Next, we tested whether the low dimensional structure derived from spontaneous activity generalized to held-out arousal transitions that had not been part of the PCA calculation. Projecting this independent PCs onto spontaneous and evoked responses showed that these components reconstructed a comparable fraction of variance at the response peak in both cases (26.2% for spontaneous vs. 27.8% for evoked arousal; **Fig. 4e)**, consistent with a shared latent structure underlying both conditions. Both PCl cart and PC2subcort were robustly recruited during arousal transitions in the same temporal ordering, with PC2subcort consistently preceding PCl ^cort^ **(Fig. 4f)**. The PCs also captured the timescale differences between conditions, with broader responses during spontaneous arousal and sharper responses during evoked arousal. Furthermore, arousal transitions traced similarly cyclical trajectories within the bidimensional subspace spanned by PCl^cort^ andPC2s^ubcort^ **(Fig. 4g)**, further supporting the generalizability of the arousal-related low-dimensional dynamics. Lastly, to test whether these results extended beyond linear methods, we applied CEBRAGS, a nonlinear embedding approach, to stimulus-free fUS data paired with pupil size **(Methods)**. When trained exclusively on spontaneous activity, the resulting latent space predicted one third of pupil responses variance on average during both spontaneous and evoked arousal transitions, outperforming temporally shuffled and session-shuffled controls **(Supplementary Fig. 8)**. Together, these analyses confirm that spontaneous and evoked arousal transitions share a common low-dimensional latent structure that is both recoverable from resting state and predictive of fUSi and pupillary responses across the two arousal conditions, consistent with the view that both spontaneous and evoked responses reflect a common arousal state.

**Figure 4.**
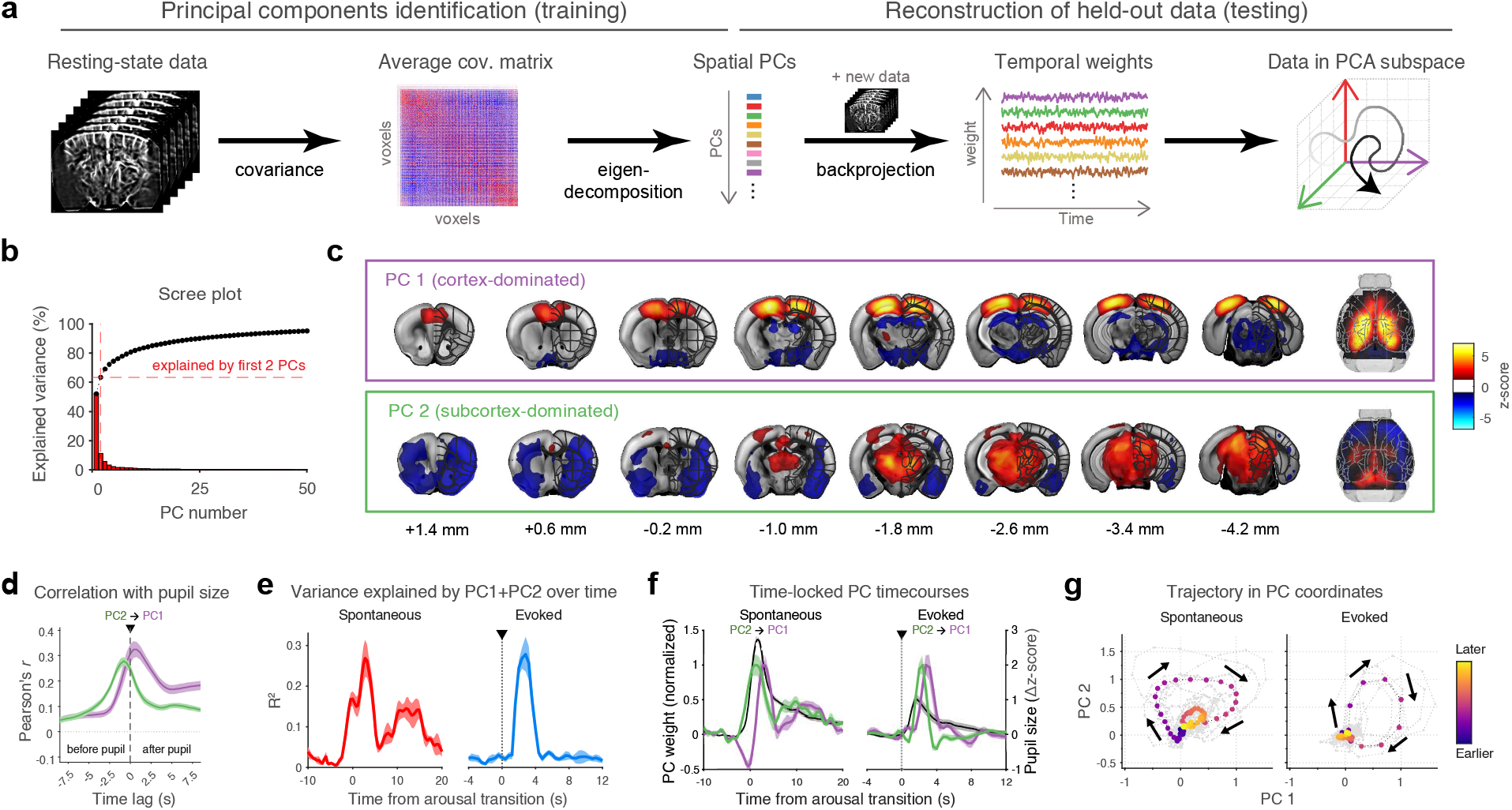
Principal component analysis captures the shared brainstem-to-cortex gradient of activation around spontaneous and evoked arousal events. a. Schematic of the group PCA approach: session-level covariance matrices are extracted from each stimulus-free recording (*n* = 45 sessions, *N* = 8 mice) and averaged. Eigendecomposition of this group-level covariance matrix returns the spatial principal components (PCs) that best explain spatial variance across all sessions. The group-level PCs are back-projected onto new session’s full fUS data, comprising both stimulus-free (*n* = 79 sessions, *N* = 7 mice) and air puffs *(n* = 21 sessions, *N* = 7 mice), to obtain temporal weights. These temporal weights can be used to time-locked analyses, but also reorganized into a low-dimensional manifold to summarize whole-brain trajectories over time. **b.** Scree plot of the normalized eigenvalues, corresponding to explained variance, of the 50 first group PCs. **c**. Left: coronal slices showing the spatial weights of the first two PCs, thresholded for absolute z-score > 1.0. Right: cortical view of the same maps. **d**. Average cross-correlation between the first two PCs and the pupil trace (n = 105 sessions, N = 15 mice), highlighting that PCl and PC2 follow pupil fluctuations with different time shifts. e. Variance explained by the combination of PCl and PC2 over time for spontaneous (*left)* and evoked (*right)* arousal. **f**. Time-locked temporal weights of arousal-related components for spontaneous (*left)* and evoked (*right)* arousal. **g**. Brain activity temporal trajectory in PCAspace for spontaneous *(left)* and evoked *(right)* arousal.

### Arousal wave is preserved during tonic manipulations of the locus coeruleus

The association between arousal state and the locus coeruleus-noradrenaline (LC-NA) system is well-established ^2,3^, with LC activity increasing in response to both spontaneous ^38-40^ and evoked^41-43^ arousal. However, while tonic LC activations have been shown to broadly modulate brain-wide signals^47-49,^ the causal relationship between LC activity - leading to NA release across the brain - and arousal-dependent brain-wide dynamics has not yet been established. To determine the impact of tonic NA modulation on fUSi activity, we expressed excitatory (ChR2) or inhibitory (stGtACR2) opsins in the LC and bilaterally implanted optic fibers behind the COMBO window **(Fig. 5a)**. Note that the space constraints imposed by the fibers forced us to use a shorter cranial window relative to previous experiments, limiting fUSi coverage of caudal regions **(Fig. 5b)**. We then applied sustained (60s) bidirectional LC manipulations (5 Hz for activation and control, continuous for inhibition) to induce prolonged changes in NA tone. Compared to the control, LC optogenetic activation produced immediate and significant pupil dilation that persisted throughout stimulation and rapidly returned to baseline afterwards. By contrast, sustained LC optogenetic silencing caused a delayed shift towards pupil constriction that however did not reach statistical significance, likely due to a physiological floor effect **(Fig. 5c,d)**. At the whole-brain level, neither manipulation elicited the previously identified arousal wave. Rather, optogenetic stimulation induced sign-consistent shifts in fUSi baseline signal during the entire duration of the manipulation period, namely decreases during LC activation and increases during LC inhibition **(Fig. 5e,f)**. These fUSi signal shifts were widespread and spatially structured, but different from the spatiotemporal organization of the arousal wave. Qualitatively, LC manipulation effects were spatially prominent in dorsal and frontal brain regions, consistent with prior reports^48^, but weak or absent in striatum and thalamus **(Fig. 5g)**. Further exploration with higher-resolution 2D-fUSi suggested further spatially heterogeneous effects of LC opto-activation near the caudal edge of the imaging window, including activation of thalamic regions, **(Supplementary Fig**. 9).

**Figure 5.**
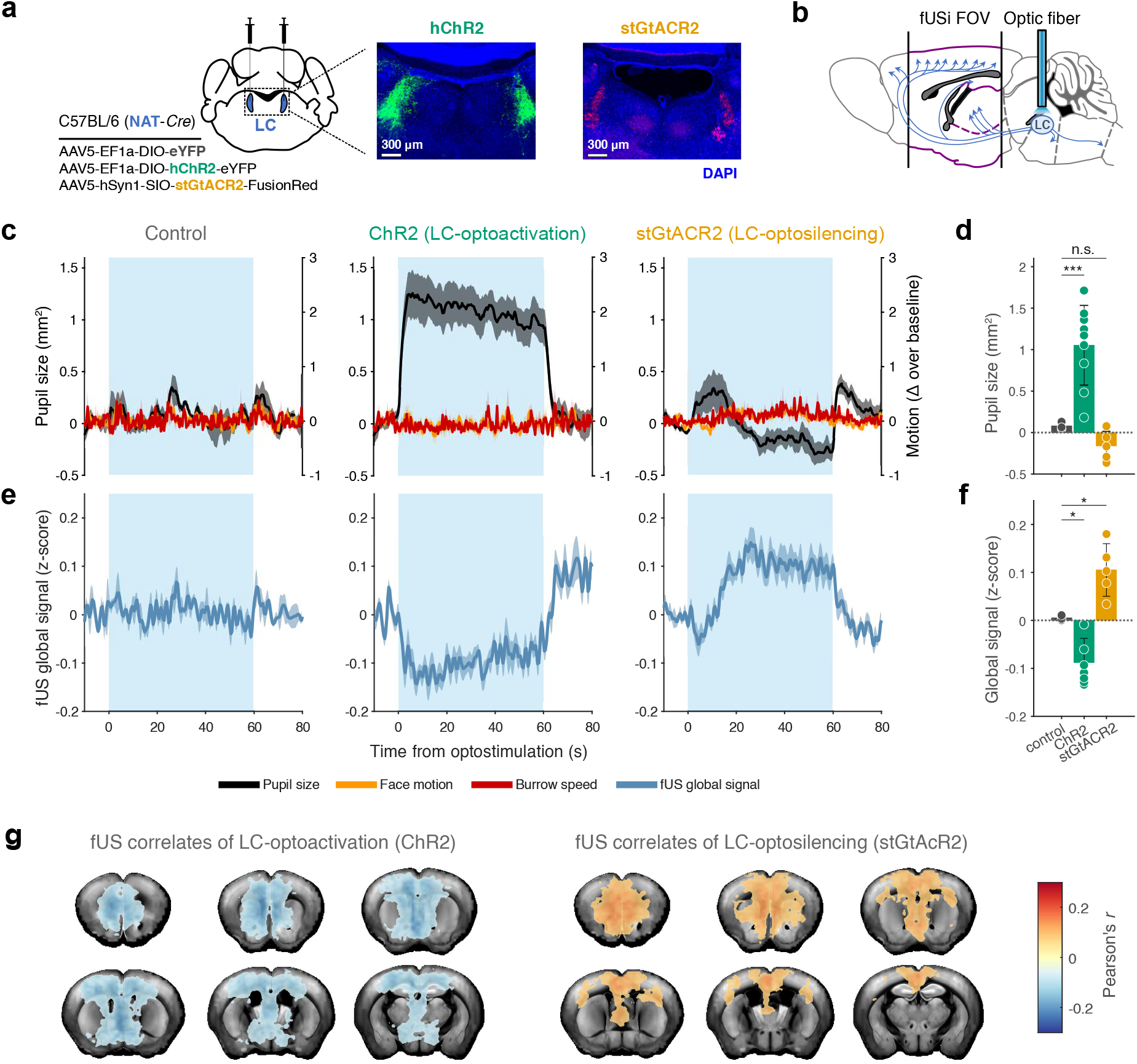
Noradrenergic tone manipulations bidirectionally affect brain perfusion. **a.** Diagram of viral injection on *NAT-Cre* animals and example confocal microscopy images showing opsin expressions for activation and inhibition cohorts. **b**. Schematics of the experimental setup, highlighting optic fiber position and shorter fUSi field-of-view. **c**. Average optostimulation-locked behavioral traces, for control *(left)*, optogenetic activation *(center)* and optogenetic inhibition *(right)* cohorts. **d**. Bar plots representing average pupil size during the last 30 s of sustained optogenetic stimulation across animals. **e**. Average optostimulation-locked global signal trace, for control *(left)*, optogenetic activation *(center)* and optogenetic inhibition (*right)* cohorts. **f**. Bar plots representing average global signal during the last 30 s of sustained optogenetic stimulation across animals. **g**. Brain-wide correlation maps of the optogenetic stimulation vector with voxelwise fUS signal (FDR<0.15), for optogenetic activation *(left; n* = 9 sessions, *N=* 9 mice, 10 cycles of 60-s ON/OFF per session) and optogenetic inhibition *(right* ; *n* = 15 sessions, *N* = 5 mice, 10 cycles of 60-s ON / OFF per session) cohorts. * = p-value < 0.05. *** = p-value < 0.001.

To further investigate the effects of LC manipulations on arousal dynamics, we sought to determine whether these tonic NA-induced effects altered the previously characterized arousal wave. Notably, although reduced in magnitude, spontaneous pupil events continued to occur during sustained LC activation and inhibition, together with fUSi responses whose spatial structure paralleled those of the arousal wave in control animals **(Fig. 6a-c, Supplementary Fig. lOa,b)**. In a complementary analysis, we examined the correlation maps between fUSi signals and face motion, another arousal proxy that is not influenced by the optostimulation itself, unlike pupil size **(Fig. 5c)**. Correlation maps remained stable across cohorts both during and outside optostimulation periods (Supplementary Fig. lOc,d), likewise indicating a lack of effect of LC manipulations over spontaneous arousal correlates. Next, to assess evoked arousal, we delivered air puffs during prolonged (5 min) LC manipulations **(Fig. 6d)**. In both activation and inhibition cohorts, air puffs elicited fUS arousal responses whose spatial extent closely matched controls and whose magnitude was only minimally affected **(Fig. 6e,f)**. Together, these results support that the transient, spatially structured fUSi dynamics associated with arousal transitions are preserved despite sustained bidirectional modulation of LC activity, suggesting that noradrenergic tone is not the primary driver of these dynamics.

**Figure 6.**
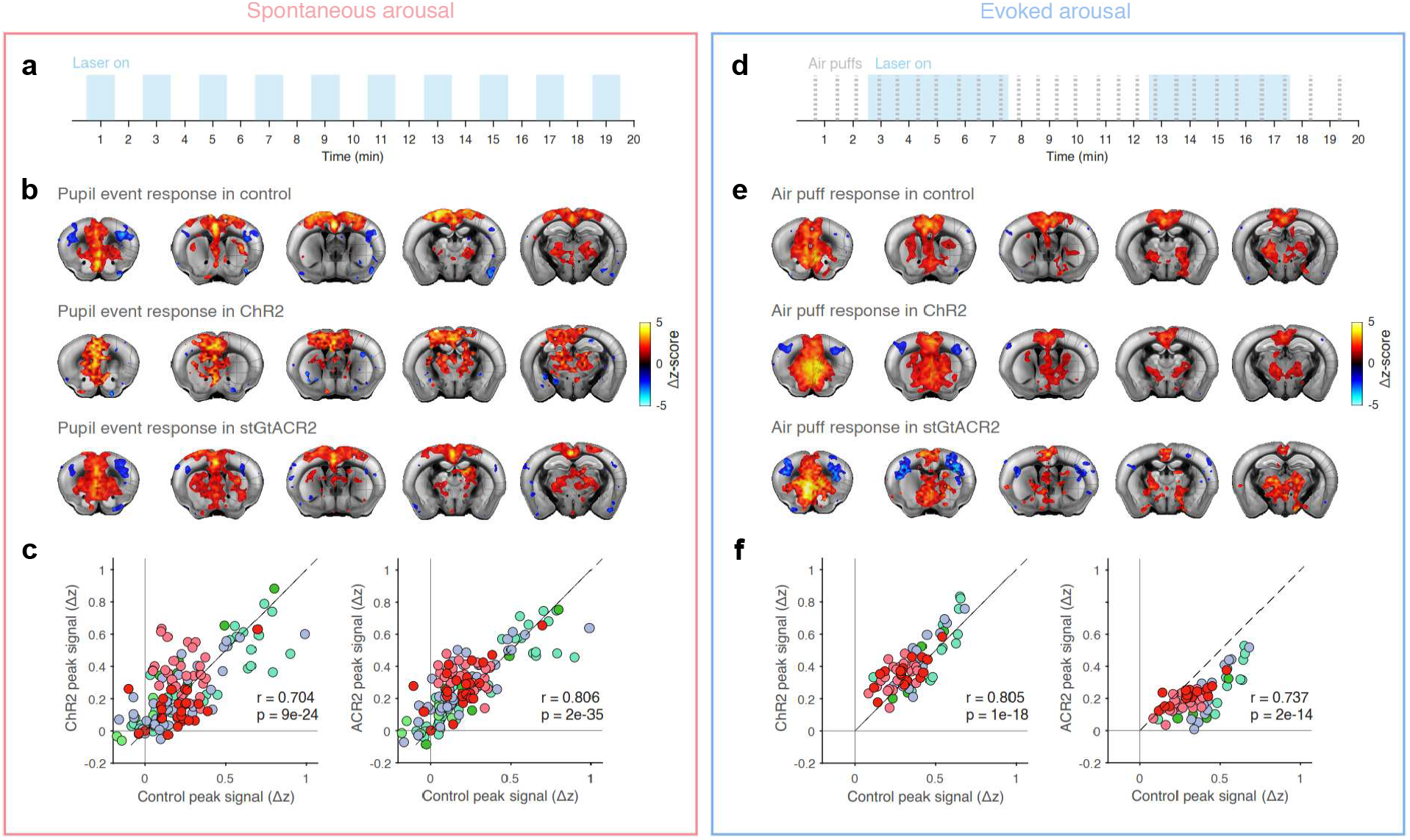
Manipulation of noradrenergic tone does not impair the arousal wave. **a.** Experimental paradigm for the stimulus-free optogenetic manipulations. **b**. Normalized (z-scored) average brain-wide tUS response amplitude for spontaneous pupil events during concurrent opto-manipulation for all three mice cohorts, overlaid with finer brain regions, thresholded for absolute z-score > 1.3. c. Scatterplots comparing each brain region’s pupil event response amplitude for the control cohort and the ChR2 (*left)* or ACR2 (*right)* cohort. Regions are defined according to the Allen Brain Atlas Common Coordinate Framework. The black dashed line indicates 1:1 correspondence. **d-f**. As **a-c**, but for air puff recordings during concurrent optogenetic manipulations

## Discussion

Arousal strongly modulates neural activity across the brain ^4,5^. While many studies have reported differential arousal effects across multiple brain regions^13-16,18,20,21^, a simultaneous characterization of arousal dynamics throughout the whole brain remains incomplete. In this study, we combine fUSi and behavioral monitoring in awake, head-fixed mice to investigate the brain’s spatiotemporal response to spontaneous and sensory-evoked arousal events. Our findings reveal a large-scale hemodynamic “arousal wave” following a subcortex-to-cortex propagation gradient across arousal conditions. The first two principal components of “resting-state” brain signals matched the spatiotemporal structure of this wave, confirming its relevance in ongoing neural activity. Moreover, we could predict both spontaneous and evoked arousal responses in test data using these components, further supporting a shared latent arousal structure. Notably, despite the long established link between noradrenaline signaling and arousal ^2,3^, the arousal wave persisted under bidirectional optogenetic manipulations of the LC, suggesting that other neuromodulatory mechanisms may play a more important role during awake arousal fluctuations.

From a technical perspective, we offer a unique characterization of arousal correlates during wakefulness that combines simultaneous whole-brain scope with behavioral monitoring in mice. Prior studies addressing differential effects of arousal in awake rodents have typically relied on electrophysiology^20^ or widefield fluorescence imaging^15,16,18,^ which have limited spatial coverage. Alternatively, rodent fMRI studies of arousal have mainly been conducted under anesthesia ^21-23,^ restricting the range of arousal states that can be accessed. Volumetric fUSi overcomes these limitations by providing simultaneous cortical and subcortical coverage during wakefulness and supporting diverse experimental configurations. In combination with a large COMBO cranial window^54^, we accessed most brain regions between the olfactory bulb and the cerebellum, excluding only lateral areas such as temporal, insular and piriform cortices.

Spatially, pupil-linked arousal effects were widespread yet heterogeneous in terms of effect size across the brain. The cortical pattern observed with fUSi closely matches previous widefield fluorescence studies, which have shown that medial motor, primary somatosensory, and anterior visual cortices strongly correlate with behavioral proxies of arousal ^14,16-18,59.^ These areas also respond to salient stimuli across sensory modalities ^10,25,29,50,61^, supporting the notion of a generalized arousal effect independent of input source. Crucially, fUSi extends and contextualizes these activations beyond the cortex. The subcortical regions revealed by fUSi include nodes previously linked to fluctuations in vigilance and pupil size^62-65^ or shown to respond to salient stimuli^25,66^, which strengthens the link to arousal. Moreover, the overall 3D pattern also resembles pupil correlates maps from fMRI studies under anesthesia - albeit with an inverted sign ^21-23^. The negative sign in fMRI correlates may reflect differences in pupil dynamics under anesthesia, such as has been observed under sleep ^67^, although the lack of comparable fMRI studies in awake rodents prevents confirming this hypothesis.

Besides spatial heterogeneity, our findings reveal that arousal courses as a structured process over time. Rather than a synchronous, global modulation, both spontaneous and evoked arousal transitions engaged a consistent wave of activation that propagated from subcortical to cortical regions along a rostrocaudal and dorsoventral gradient. This spatiotemporal ordering was reproducible across animals and validated in an independent fiber photometry dataset, supporting its robustness across measurement modalities. Moreover, such a progression matches prior reports of regional timing differences during spontaneous ^18-20-24^ and evoked ^68^ arousal, as well as the propagation of low-frequency waves of activity accompanying awakenings in humans^69^. In contrast, a recent 2D-fUSi study did not detect large temporal offsets around spontaneous whisking events in the absence of locomotion ^24^, possibly due to the smaller amplitude of the arousal fluctuations examined. This temporal evolution may therefore reflect a fundamental, time-evolving neural process along bottom-up arousal pathways, including the midbrain-to-cortex projections from the reticular system ^36^, that sequentially prepares the brain for interaction with the external world during large-scale arousal events.

Our PCA analysis further recapitulated the arousal temporal structure in an independent stimulus-free dataset via two dominant components: an early subcortical component and a later cortical component. Prior PCA-based studies of arousal have largely focused on cortical recordings, typically identifying a single dominant arousal component ^19-56-57^. In contrast, our whole-brain approach reveals the subcortex as a distinct contributor to arousal-related low-dimensional structure. When used to reconstruct held-out fUS data, this bidimensional representation explained more than a quarter of the variance in both spontaneous and evoked arousal responses, thus quantifying the shared latent structure across arousal conditions. This fraction of explained variance is in line with former studies of more restricted spatial scope^16,17,19,20^, suggesting a consistent upper bound for the common arousal contribution to session-level brain activity fluctuations. A similar result was further obtained from the non-linear algorithm CEBRA, which could predict both spontaneous and evoked pupil responses on a testing set. These observations thus align with the notion of “generalized arousal”: a common latent brain-body state that underlies different physiological and behavioral phenomena - such as our spontaneous and evoked conditions - in a context-dependent manner ^1,55,55,70^. Notably, the arousal wave described here also resembles large-scale co-activation patterns from fMRI studies linked to physiological variables ^71^, global signal topography ^72^, and default-mode network ^25,73^, consistent with a core role in brain function.

Mechanistically, the LC-NA system is often considered a core substrate of arousal ^2,3,74^ .However, our optogenetic LC manipulations did not reproduce the arousal wave, but instead caused widespread decreases (for optogenetic activation) or increases (for optogenetic silencing) in fUS signal amplitude, consistent with previous pharmacogenetic and optogenetic fMRI studies^46-48.^ Further, these modulations appeared spatially structured, with no significant effect in the striatum and the thalamus. While striatum does not receive profuse LC innervation, the lack of thalamic effect likely stems from low signal-to-noise ratio, in turn caused by the shorter fUSi field-of-view imposed by the optic fiber placement over the LC. Supporting this interpretation, single-plane fUSi recordings in the mice with most caudal cranial windows identified thalamic activations in response to LC stimulation, as reported in prior publications^46-47.^ Regardless of these effects, both spontaneous and evoked arousal events retained the characteristic fUS arousal pattern during sustained bidirectional LC manipulations. Of particular interest, LC silencing did not abolish the arousal wave, suggesting its progression to be independent of tonic noradrenaline. While it is still possible that our inhibition was not absolute due to incomplete opsin expression, recent work in the hippocampus has identified different neuronal effects between optogenetic LC stimulation and natural arousal ^75^, suggesting a more complex modulation of the latter. In that regard, a recent fl\1RI study showed that cholinergic optostimulation evokes a whole-brain pattern closely resembling the one presented here ^76^, thus positioning acetylcholine - which is also strongly associated with the behavioral proxies of arousal ^40,59,77-79^ and the brain response to salient stimuli ^80,81^ – as a likely contributor to the physiological arousal wave.

Alternative interpretations of our results could invoke sensory, motor or hemodynamic confounds, all of which are intertwined with arousal-related fUS signal. Several controls nevertheless constrain these explanations. Regarding a sensory component, our anesthesia experiments confirmed that deep sedation abolishes much of the air puff-evoked fUS response, sparing only well-known sensory-integration hubs, indicating that the widespread pattern cannot be attributed to sensory responses alone. Light-driven pupil changes likewise failed to reproduce the whole-brain correlates of arousal-driven pupil fluctuations, arguing against purely visual or pupillary explanations. Motor contributions were minimized by stimulus calibration and further dissociated from arousal via partial correlation analyses, although residual motor contributions cannot be fully excluded. A more fundamental limitation concerns the hemodynamic nature of fUSi, which, like fl\1RI, measures blood volume changes rather than neural activity directly. While previous studies have shown that fUS signals reliably track bulk neural activity in the low frequencies ^52,82,83^ - the frequency range of arousal fluctuations ^7^ -, other works have identified differences in hemodynamic coupling across arousal states ^67,84^ and across arousal-responsive neural populations ^24^. Given that noradrenaline directly and indirectly affects the vascular tone^85-89^, it is possible that LC optogenetic manipulations likewise induce neurovascular decoupling. Importantly, a recent preprint reported global inhibition of hippocampal neurons in response to LC stimulation ^75^, suggesting that large mismatches between the fUSi signal decreases and actual neural activity are unlikely.

In summary, our study uncovered a stereotyped brain-wide hemodynamic wave associated with both spontaneous and evoked arousal events. This wave accounts for a substantial fraction of resting-state brain variance, indicating a major contribution to ongoing brain dynamics. Such a wave is compatible with a generalized arousal latent state that regulates brain function regardless of whether arousal arises from an internal state change or a salient sensory event. Notably, noradrenergic tone manipulations did not prevent expression of the arousal wave, suggesting a role of other neuromodulatory mechanisms, such as acetylcholine. Elucidating these mechanisms and their neuromodulatory interplay thus represents a natural direction for future work.

## Methods

### Mice

Male and female C57BL/6 mice (>8 weeks of age, weighted 15-30 g) were used in this study. For optostimulation experiments, a transgenic mouse line expressing Cre recombinase under the control of the noradrenaline transporter gene (NAT, Tg(Slc6a2-cre)FV319Gsat) from the Max Planck Institute of Psychiatry internal colony was used; the rest of the experiments employed wild-type mice from the internal colonies of the Max Planck Institute of Biological Intelligence and the University Medical Center of Goettingen. All animals were group-housed (3-5 individuals/cage) in the standard laboratory condition (temperature: 23 ± 1 °C; humidity: 50-70%) under a 12h reversed light/dark cycle, with food and water ad libitum. All behavioral experiments were performed during the dark cycle. All animal procedures were performed in accordance with the institutional guidelines of the Max Planck Society, the government of Upper Bavaria (Regierung von Oberbayern), and the Lower Saxony State Office for Consumer Protection and Food Safety (Niedersachsisches Landesamt for Verbraucherschutz und Lebensmittelsicherheit).

### Surgical procedures

#### Craniotomy

All surgical procedures were conducted under standard aseptic conditions. Mice were anesthetized with a subcutaneous injection of a fentanyl (0.05 mg/kg), medetomidine (0.5 mg/kg), and midazolam (5 mg/kg) (FMM) cocktail and implanted with a COMBO window as previously described ^54^. In brief, the scalp was shaved and disinfected with iodine, and an eye ointment (Bayer, Bepanthen) was applied over the eyes to prevent corneal dehydration during surgery. Mice were then secured using a bite bar and placed on top of a heating mat (Supertech) to maintain a body temperature of 37°C. The scalp was cut and a dental drill with a 0.5 mm burr (Hager & Meisinger) was used to delineate a large cranial window into the skull, extending from Bregma +2.00 mm AP to Bregma -4.50 mm AP, and along the full width of the dorsal skull. The resulting bone island was carefully removed with a forceps keeping the dura intact. A pre-prepared 3D-printed COMBO implant was then attached to the remaining bone using cyanoacrylate glue (Pattex) and sealed with dental cement (Super-Bond). After surgery, anesthesia was reversed with a subcutaneous injection of a cocktail of flumazenil (0.5 mg/kg) and atipamezole (2.5 mg/kg). Buprenorphine (0.1 mg/kg) was injected subcutaneously for post-surgical analgesia. Mice were then placed in a warm cage and monitored until they woke up. The health of the animals was evaluated by post-operative checks during the five following days and buprenorphine (0.1 mg/kg) was administered subcutaneously when needed. After seven days of recovery, animals were again anesthetized with FMM as previously described and a stainless steel head-plate was attached to the window frame to enable later head-fixation, after which anesthesia was reversed with flumazenil-atipamezole. Animals were left to recover for seven additional days before the first imaging session. During the fourteen total recovery days, mice were carefully handled to habituate them to the experimenter.

### Vims injection and fiber implantation

For animals used in optogenetic experiments, virus injection and fiber implantation were performed prior to the craniotomy in the same surgery. Animals were anesthetized using the FMM cocktail previously described and secured on a stereotaxic frame (Stoelting) with ears and teeth fixed. After exposing the skull and drilling a small hole over the target site, mice were injected bilaterally in the locus coeruleus (Coordinates: AP: -5.4 mm, ML: ±1.0 mm, DV: -3.8 mm, relative to Bregma) with 500 nL of an AAV construct expressing channelrhodopsin (AAV5-EFla-DIO-hChR2(H134R)-eYFP; University of North Carolina vector core, #4313), 325nl nL of an AAV construct expressing anion-conducting channelrhodopsin (AAV5-hSynl-SIO-stGtACR2-FusionRed; AddGene, #105677) or 500 nL of a control AAV construct (AAV5-EFla-DIO-eYFP; Vector Biolabs, #VB2089) using a pneumatic injector (Stoelting) and pulled capillary glass calibrated micropipettes (Sutter instrument) at a rate of 50 nL/min. After injection, the glass needle was left in place for 10 min to avoid leaking and then carefully removed. Subsequently, low-profile optical cannulas (low profile 90°, 200µm, NA=0.66; Doric Lenses) were implanted at 0.2 mm above the injection coordinates and stabilized on the skull with cyanoacrylate glue (Pattex). A shorter craniotomy was then performed, ranging from Bregma +2.00 mm AP to Bregma -2.50 mm AP, and a fiber-compatible COMBO window^54^ was placed on the remaining skull. Finally, both the fibers and the window frame were fixed with dental cement (Super-Bond) and the surgery ended as described above.

Animals used in fiber photometry recordings were injected bilaterally with 500 nL of an AAV construct expressing GCaMP (AAV9-syn-jGCaMP7f-WPRE; Addgene, #104488). Injection sites were the taenia tecta (Coordinates: AP: -1.7 mm, ML: 0.0 mm, DV: -3.5 mm, all coordinates relative to Bregma), the frontal cingulate cortex (AP: +1.2 mm, ML: ±0.5 mm, DV: -1.8 mm), the caudal cingulate cortex (AP: +0.1 mm, ML: ±0.35 mm, DV: -1.5 mm), and the somatosensory cortex (AP: -1.5 mm, ML: ±1.8 mm, DV: -0.6 mm). The procedure was the same as for the one used in optogenetic experiments, except that custom-made optical cannulas were implanted over the target site instead of low-profile ones, no craniotomy was performed afterwards, and the window frame was cemented on the intact skull to enable head-fixation.

### Behavioral monitoring

#### Virtual burrow assay

To monitor behavioral arousal while recording fUS signals, animals were head-fixed using two custom-machined clamps and placed within a virtual burrow assay (VBA)^53^. In brief, the VBA is a 3D-printed tube enclosure (the virtual burrow) mounted on near-frictionless air bushings (New Way Air Bearings) that slide on two parallel rails to constrain movement in one dimension, and tethered to a linear actuator with a fishing line to control its motion range. Inside the virtual burrow, mice heads were tilted for 30 degrees to achieve a more naturalistic posture and reduce stress. A trimmed absorbent underpad was attached to the floor of the tube, underneath the mouse, before each imaging session. The complete VBA set-up was placed inside a sound-attenuating black box over an anti-vibration table to insulate animals from external stimuli. Animals were acclimated to the VBA for 2 min after head fixation. The mice were free to go back and forth within the virtual burrow and typically entered the tube immediately and remained calm inside, egressing only during arousal bouts. The movement of the virtual burrow around the mice’s body was tracked using a laser displacement sensor focused on a 3D-printed flag that was affixed to the side of the tube enclosure. The analog voltage signal from the displacement sensor was digitized at 1 kHz using a NI-DAQ card (National Instruments, USB-6001).

#### Face and pupil recordings

The mouse face, from ear base to snout, was recorded at 30 frames/s with a monochromatic camera (Flir, BFS-U3-13Y3M-C), positioned at the right side of the mouse head, using the SpinView software (FLIR). Two infrared light-emitting diode (LED) arrays (875 mm, Kemo Electronic, M120) were used to enhance the contrast of the pupil and whiskers images, and a 720 nm filter (Hoya, R72 E49) was attached to the camera to remove non-IR light. Mice were weakly illuminated from the front with a small white LED light (Varta) to keep the pupil relatively constricted in non-aroused conditions. To synchronize video frames with other data, videographic recordings were initiated with a transistor-transistor logic (TTL) trigger from a master NI-DAQ card (National Instruments, USB-6211) controlled with custom software written in MATLAB (MathWorks).

### Experimental manipulations

#### Sensory stimulation

Each sensory stimulation session lasted for 16 min. During the first 6 minutes no stimulation was present in order to record spontaneous behavioral transitions from the mice. During the next 10 minutes, somatosensory or auditory stimuli were presented in a pseudorandomized way, with average inter-stimuli intervals of 30 s. In the case of anesthetized sessions, mice were injected with an FMM cocktail prior to the imaging session and recorded for 30 minutes, after which the anesthesia was reversed with a flumazenil-atipamezole cocktail.

Somatosensory stimuli were brief (50 ms) or sustained (10 s) weak air puffs (air pressure 30 psi) delivered towards the left side of the face via a blunt 18G needle tip placed 20 cm away from the animal and connected to a silicone tube (Cole-Palmer) that passed through a normally-closed 12 V pinch valve (NResearch). The valve opening was controlled by the master NI-DAQ system via a valve driver (NResearch, CoolDrive®). Auditory stimuli were 0.5 s bouts of broadband Gaussian noise (central frequency: 14 kHz, FWHM: 7 kHz) delivered at 70 dB via a USB speaker (RS Components) placed 20 cm away in front of the animal.

The audio signal and air-puff valve triggers were generated with the same master MATLAB script that initiated the video and fUSi acquisitions (MATLAB *sound, daq, start, read)*. The timestamps of the stimuli were stored for later analysis.

#### Light-driven pupil fluctuations

Mice were exposed to a green mounted LED (Thorlabs) connected to a modulable LED driver (Thorlabs), which in turn was controlled by the master NI-DAQ system. Imaging sessions started with 6 minutes of constant illumination to enable paired analysis within the same cohort. Immediately afterwards, the driver induced a “saw” pattern of illumination where the LED light ramped up and down for the next 10 minutes, with 30 between maximum and minimum illumination, therefore 1 minute between two peaks or two troughs. The slow ramping length was specifically chosen to minimize phasic stimulation of the visual pathways. The light intensity was calibrated for each mouse so that maximum and minimum light corresponded to minimum and maximum pupil dilation, respectively.

#### Optogenetic stimulation

All optogenetic stimulation experiments took place 4 weeks after virus injection to allow for sufficient expression. The implanted cannulas received light from a 473 nm laser (CNI Laser) via a 200 µm diameter, 0.22 numerical aperture optic patch cord (Doric Lenses Inc.). The patch cord fibers and the animal cannulas were attached using bronze mating sleeves (Thorlabs), which were carefully covered with black plasticine to prevent light leak. Before each session, the laser power was adjusted to 10 mW (control and optostimulation) or 4 mW (optosilencing) at the tip of each cord fiber. Optostimulation was delivered as trains of 10-ms-long light pulses at 5 Hz, whereas optosilencing involved continuous light emission. In both cases, optogenetic manipulation spanned two different total lengths: 60 s (60 s inter-stimuli time) for spontaneous recordings, and 300 s (300 sinter-stimuli time) for concurrent air puff stimulation. Total recording time was 20 minutes per session. The manipulation paradigms were controlled by a pre-programmed pulse generator (Doric, OTPG_8).

### fUS acquisition

fUS imaging data was acquired using a 32 x 32 channel matrix probe (15 MHz, 1024 total elements, spatial resolution: 220 x 280 x 175 µm^3^, Vermon) attached to a 256-channel workstation (Vantage 256, Verasonics) via a 4x multiplexer (Verasonics). At the beginning of each imaging session, before placing the ultrasound probe, acoustic gel (Sonosid) was applied on the cranial window for ultrasound coupling. The acoustic gel was frequently centrifuged to avoid air bubbles and kept at room temperature. The matrix probe was positioned onto the gel to encompass the full width and length of the cranial window using a three-way translation stage, 3 mm above the plastic surface of the cranial window. The probe position was verified by acquiring single Doppler images until achieving optimal coverage and alignment per session.

The beamforming and plane-emission sequences were controlled by a custom volumetric fUSI acquisition module (AUTC). In brief, a single compound ultrasound image was generated from the summation of the reconstructed echoes of plane wave emissions at −4.5, −3, −1.5, 0, 1.5, 3, and 4.5 degrees. Blocks of 160 compound ultrasound images, acquired at a pulse repetition frequency of 400 Hz, were stacked to create a single power Doppler image. Clutter filtering was performed in real time, whereby the compound ultrasound stack was decomposed using singular value decomposition and the first 20% of singular vectors were excluded to remove artifacts due to brain movement ^90^. The power Doppler image was then created via incoherent average, by averaging the squared complex-valued compound images. This procedure produced a single power Doppler image every ∼400 ms. The beginning of the fUS recording was synchronized to the behavioral data acquisition and the stimuli delivery through a master computer.

To manually register the fUS data to the Allen Brain Atlas space, a high-resolution 3D anatomical scan of the brain microvasculature was acquired at the end of the first imaging session of each mouse. The high-resolution Doppler was obtained as described above, but used plane wave emissions at −9, −6, −3, −1, 1, 3, 6, and 9 degrees to increase resolution.

To obtain detailed images of hemodynamic propagation, 2D-fUS was applied to specific brain slices. In this case, the process was identical to that of 3D-fUS except that it employed a 1 x 128 channel linear probe (15 MHz, spatial resolution: 100 x 113 µm2, Vermon) applying plane waves at −10, −8, −6, −4, −2, 0, 2, 4, 6, 8, and 10 degrees. Power Doppler compounding used blocks of 200 compound ultrasound images and took ∼200 ms to produce each image.

### Fiber photometry recording

All photometric recordings began at least 4 weeks after virus injection to allow for sufficient expression. Fiber photometry was performed using a commercially available system (Doric Lenses). To detect mesoscopic GCaMP fluorescence, two LEDs were used to deliver interleaved excitation light: a 465 nm LED for recording calcium-dependent fluorescent signal and a 405 nm LED for calcium-independent (isosbestic) control fluorescent signals. Power in the two LEDs was adjusted to achieve comparable intensity of emitted fluorescent light for both channels. The elicited fluorescence was acquired via two pre-bleached 200 µm diameter, 0.57 numerical aperture optic patch cords (Doric Lenses), each one connected to a 4-ports fluorescence minicube (Doric Lenses). Inside the minicubes, the fluorescence signals were collected by integrated photodetectors, amplified and digitized at a sampling rate of 40 frames/s.

### Histology

At the end of the experimental protocol, mice used for optogenetic or fiber photometry experiments were deeply anesthetized with an intraperitoneal injection of ketamine (300 mg/kg) and xylazine (30 mg/kg) and transcardially perfused with cold 0.1 M phosphate-buffered saline (PBS, Life Technologies) followed by 4% paraformaldehyde (PFA, Thermo Scientific) in PBS. Brains were postfixed by immersion in 4% PFA for 24 hat 4°C. Following a rinse in PBS, brains from the fiber photometry cohort were stored in 30% sucrose (Sigma-Aldrich) in PBS solution for 48 h for cryoprotection and then frozen in NEG-50 freezing medium (Fisher Scientific) inside cryo-molds (Sakura), after which they were sectioned into 50 mm coronal slices using a cryostat (Leica) and directly mounted on glass slides (Thermo Fisher Scientific). Brains from the optogenetic cohort were embedded in 4% agarose (Sigma-Aldrich) and sectioned into 70 µm coronal slices using a vibratome (Leica), after which the floating sections were stained with DAPI (Invitrogen™, NucBlue™(Hoechst 33342)) and mounted on glass slides. Images were acquired using an epifluorescent light microscope (Nikon, Eclipse) with a 2x and l0x objective, or a confocal laser-scanning microscope (Leica, Thunder) with a l0x objective. LAS X (Leica Microsystems) and ImageJ were used to process the acquired images.

### Data analysis

All subsequent analyses were performed using custom scripts in MATLAB R2021a (Math Works).

#### Behavioral data preprocessing

Virtual burrow speed was computed as the differential of the low-pass filtered (< 20 Hz) laser displacement trace. Pupil diameter and facial movements were extracted from face videos using the FaceMap software^91^ (www.github.com/MouseLand/FaceMap) and smoothed with a Savitzky-Golay filter (MATLAB *sgolayfilt)* of order 10 and length 1 s. Blinks were identified as negative peaks (MATLAB *findpeaks)* in the size and change-rate traces of the pupil signal, padded by 0.4 s and linearly interpolated (MATLAB *interp1*). All three signals were appropriately downsampled (MATLAB *interpl)* to match the fUSi or fiber photometry sampling rates (2.5 and 30 frames/s, respectively).

#### Fiber photometry preprocessing

Signals from both fiber photometry channels (405- and 465-nm) were first smoothed with a 2.5 s moving median filter (MATLAB *movmedian)* to remove high-frequency noise. Next, to correct for movement artifacts and photobleaching of the fluorescent signal, the 405-nm signal was fitted to the 465-nm signal by using a least-squares linear fit (MATLAB *polyfit)*, and the *ΔF*/ *F* signal was calculated as the percentage of increase of the 465-nm signal over the fitted 405-nm signal:

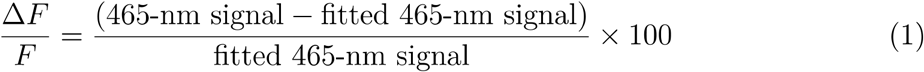

The first 15 s of each *ΔF* / *F* signal were removed to eliminate initial bleaching artifacts and achieve a steady signal, and the resulting traces were downsampled to 30 frames/s (MATLAB *interp1*).

#### JUS signal preprocessing

For each mouse, a high-resolution 3D image of the brain microvasculature was manually registered to the reference mouse brain atlas from the Allen Brain Institute (https://atlas.brain-map.org/) according to anatomical landmarks in order to obtain a mouse-specific transformation matrix (*M*_4×4_). Registration to atlas included translation, rotations, and scaling steps, resulting in a 4 x 4 affine transform matrix. Additionally, reference images from all the sessions of each mouse, calculated as the average of the 100 fUS frames with the lowest motion, were automatically registered via Mattes mutual information maximization (Matlab *imregtform)* to this high-resolution session in order to get a session-specific transformation matrix (*S*_4×4_). Because inter-session variability within animals is smaller than inter-animal variability, no scaling step was used for the automatic registration, resulting in 4 x 4 rigid transform matrices. Both matrices were combined via matrix multiplication to obtain session-to-atlas transform matrices (*T*_4×4_ = *S*_4×4_ × *M*_4×4_) that could be applied during the preprocessing stage. No motion correction was performed between timepoints, but rather the same transform matrix was used for all frames.

Each session’s fUS time series were then preprocessed on a voxel-by-voxel basis. First, data was temporally interpolated to correct for timing differences in fUS acquisition and to obtain a consistent frame rate of 2.5 Hz (MATLAB *interp1*). Then, the aforementioned transformation matrices were applied to the data to register it to the common space template (MATLAB *imwarp)*. A logical mask defining brain vs. non-brain voxels was manually drawn in the aforementioned high-resolution fUS image using the ITK-SNAP software (http://www.itksnap.org/). Next, to remove nonspecific movement artifacts and other non-neuronal sources of noise, the top 1.5% principal component of the non-brain voxels fUS signal, i.e., skull and skin (MATLAB *sud)*, was linearly regressed from the whole signal, and a low-pass FIR filter with cutoff frequency of 0.5 Hz ^92^ was subsequently applied (MATLAB *firls, filtfilt*). Lastly, the relative change in power Doppler signal was calculated by first subtracting the average signal from each time point and then dividing the result by the standard deviation (MATLAB *zscore)*.

#### Power spectra and coherence analyses

Spectral power was calculated via the fast Fourier transform of the studied timeseries (MATLAB *fft)*. Analyses of coherence between pairs of signals employed the Welch’s overlapped averaged periodogram method for magnitude-squared coherence (MATLAB *mscohere)*, calculated up to the Nyquist limit (15 Hz for behavioral and fiber photometry traces) with a 0.01 Hz resolution and a 70% overlap.

#### Cross-correlation and partial correlation

The correlation between fUS signals and behavioral or stimulus traces was considered as the maximum value from the cross-correlation between them. The cross-correlation is a measure of similarity between a vector x, and shifted (lagged) versions of vector y. Unlike Pearson’s (0-lag) correlation, it accounts for temporal differences between the studied time series. To obtain correlation maps, mass univariate cross-correlation was efficiently calculated voxel-wise as the convolution of vector x and the complex conjugate of each voxel yin the fUS signal matrix (MATLAB *conv2, conj)*. In the cases where only two traces were compared, the cross-correlation built-in function was used instead (MATLAB *xcorr)*. Cross-correlation values were normalized so that autocorrelations at zero lag equaled one. To obtain a more precise estimation of the cross-correlation peaks, the 3 maximum values were parabolically interpolated before extracting them ^93^. Peak correlation values were Fisher-transformed to create approximately normal distributions (MATLAB *atanh)* and averaged across sessions to obtain individual mouse-level results. All statistical tests were performed on the mouse level, after which the individual maps were averaged to obtain the group results and the Fisher transformation was reversed for display (MATLAB *tanh)*.

The lag until the maximum cross-correlation is a metric of the temporal shift between two signals, so it can be used to temporally arrange responses over time; for instance, indicating which behavioral trace or which brain region activity peaks first. This metric was used in order to calculate the average lag between motion and pupil dilation, the temporal relationship between each fUS voxel and the pupil size trace, and the relative sequence of activation of the arousal regions.

For control analyses assessing the unique contributions of different regressors, partial correlation was used instead of absolute maximum cross-correlation. Partial correlation is another metric of similarity between two traces that derives from linear regression models. Unlike Pearson’s (full) correlation, partial correlation estimates covariance after regressing out effects from other control variables, and thus measures direct or unique effects (MATLAB *partialcorri)*. Partial correlation values were Fisher-transformed to create approximately normal distributions (MATLAB *atanh)* and averaged to obtain the group results, after which the Fisher transformation was reversed (MATLAB *tanh)*.

#### Arousal transitions-locked signal analysis

For both spontaneous and evoked arousal, frames with arousal transitions were identified. For spontaneous arousal transitions, pupil events were defined as instances where the pupil size increased by more than 1.5 standard deviations, identified as local maxima in the change rate of the smoothed z-scored pupil size trace (MATLAB *islocalmax, diff, sgolay, zscore)*. Event center was defined as the timepoint of maximum pupil change-rate. Events that happened less than 10 s after another were discarded. For evoked arousal, the sensory stimuli timestamps were used. In both cases, fUS and behavioral traces were extracted from 10 s before to 20 s after the transition frames. Each time-locked trace was subtracted the average value of the first 5 s of the series (from 10 s to 5 s before the transition) to correct for baseline differences and then averaged per mouse.

To identify which fUS voxels were more affected by the arousal transitions, the amplitude of each voxel response was defined as its maximum absolute value around the pupil event (from 2 s before to 5 s after the maximum pupil dilation) or after the sensory stimulation (from O s to 5 s after the stimulus). In both cases, the temporal range was chosen to encompass the full average pupil dilation. The maximum absolute value was chosen instead of the maximum value to account for potential decreases in the fUS signal. Each temporal trace was smoothed with a Savitzky-Golay filter of order 3 (MATLAB *sgolayfilt)* and parabolically interpolated before extracting their amplitudes in order to increase the temporal precision. The resulting set of amplitudes was averaged first per mouse and then across mice in order to obtain the group activity map. In addition, the time until the maximum absolute value (time-to-peak) was extracted voxel-wise and averaged per and across mice in order to obtain group delay maps.

To estimate the responses’ replicability, each mouse’s time-locked traces and voxel-wise peak amplitude maps were extracted as described above. The spatial correlation coefficient between all possible frames from all mice was calculated to obtain consistency over time across individuals. Spatial correlation was also calculated for the peak maps across mice. For the peak maps, intraclass correlation was defined as the degree of absolute agreement among measurements made on k randomly sampled subjects, as per the formula: *ICC= (MS*_*B*_*-MS*_*w*_*)/(MS*_*B*_ +*(k+* 1)*\*MS*_*w*_), where *MS*_*B*_ is the between-subject variability (mean square) and *MS*_*w*_ is the within-subject variability. In addition, to estimate the reliability of the response across mice, i.e., how similar was each individual to the average, an specificity index was calculated as (spatial correlation with a correct template) − (spatial correlation with an incorrect template) ^94^, where the correct template was the average map across mice and the incorrect template was the average of the pre-arousal baseline, representing random responses.

In order to determine the most affected brain regions, fUS voxels that were significantly active for both spontaneous and evoked arousal were parcellated according to the Allen Mouse Brain Atlas anatomical labels^95^. Each anatomical region time trace was considered to be the average between the time traces of all the voxels within that region, per mouse. Each region’s average time-to-peak was obtained from these time traces as previously described for voxels. The time traces were over-sampled to 10 Hz (MATLAB *spline)* for display, but not for analyses. To estimate the temporal sequence of activation between anatomical brain regions, the regional time traces were cross-correlated, as previously described, with the average signal of all brain voxels per mouse. To display the consensus temporal organization of the arousal response, both spontaneous and evoked regional lags were transformed into ordinal ranks, so that they could be compared across different timescales, and averaged.

#### Group PCA

Group spatial principal component analysis (PCA) was performed across all stimulus-free fUS recordings (MATLAB *eigs)*. Due to the large size of this dataset, group PCs were estimated from an average covariance matrix calculated iteratively, instead of from concatenated data. Briefly, each individual fUS scan, represented as a bidimensional time-space matrix of *t* time points and *v* voxels *(Y*_*t×v*_), was left-multiplied by its transpose to obtain its spatial covariance matrix 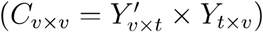. Of note, these fUS scans were z-scored to avoid small clusters of voxels (as resulting from motion artifacts) to dominate the session variance, so that these covariance matrices are formally correlation matrices. Moreover, since the first principal component always reflected the session global signal (average of all voxels) and explained most of the variance (>90%), we regressed the global signal out of all the fUS voxels prior to the covariance calculation. The resulting matrices were scaled by the number of time points t within their session to reflect estimation robustness and averaged across session. The weighting was then reversed and the individual-level covariance matrices were averaged across mice to produce a single group-level matrix. This average covariance matrix underwent sparse eigendecomposition *(W*_*m×v*_ ×*C*_*v×v*_ = *Λ*_*v×v*_ × *W*_*m×v*_, *m* < *v*), thus resulting in m spatial components *(W)* and their corresponding eigenvalues *(Λ)*, which represent explained variance after normalization.

The group spatial PCs were back-projected to each fUS session through matrix multiplication to calculate their temporal weight in each individual session 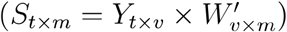. These temporal weights, equivalent to group temporal eigenvectors, quantify the similarity between each spatial PC and the fUS signal per frame, with high weights signaling frames where the spatial organization of the fUS scan resembles that of the PC. Cross-correlation between the PC temporal weights and behavioral variables, and arousal transitions-locked analysis of the PC temporal weights, were carried out as described above for the fUS signal. To estimate how much temporal variance (*R*^2^) was explained by the two arousal-related PCs, mouse-level arousal-locked fUS volumes were reconstructed from the PCs spatial and temporal weights (Ŷ _*t*×*v*_ = *S*_*t*×2_ × *W*_2×*v*_) and 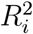; was defined as the squared spatial correlation coefficient between *Y*_*i*×*v*_ and Ŷ_*i*×*v*_ for each timepoint *i*.

#### CEBRA

The Contrastive Embedding for Behavioral and Neural Representation Analysis (CEBRA) ^58^ framework was used to map high-dimensional fUS time series into a low-dimensional latent space. CEBRA is a self-supervised framework designed to extract the latent structure of a dataset (embedding) by bringing temporally or behaviorally similar activity patterns closer together while pushing dissimilar patterns apart. Unlike the other analyses, CEBRA was implemented in Python following the instructions provided in its documentation (https://cebra.ai/docs/).

Temporally z-normalized, anatomically parcellated fUS recordings were classified into “training” and “testing” sets, each one containing non-overlapping full sessions from different mice. CEBRA-Behavior models were trained from concatenated stimulus-free sessions and concurrent pupil size, using the following parameters: model_architecture=offsetl0-model, batch_size=512, learning_rate=le^−4^, temperature_mode=auto, min_temperature=0.1, max_iterations=500, output_dimension=3, and distance=cosine. A k-nearest neighbors (kNN) algorithm was trained to decode pupil size from the resulting embedding with k selected via coarse search over the range [10, 5000] and “cosine” distance metric. Both the trained CEBRA model and the trained kNN decoder were applied to test fUS data from both stimulus-free and air puff sessions. To measure the embedding consistency, session-specific models were also obtained and the inter-session *R*^2^ scores were calculated as:

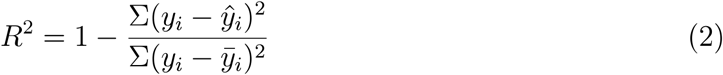

Control CEBRA models were trained using 1) pupil traces from unrelated sessions, so that the fUS-pupil correlation was lost, and 2) randomly permuted pupil traces, so that their temporal properties were lost. Finally, in order to infer the relative decoding importance of each anatomical brain region, additional models were trained based on fUS data where all anatomical parcels except those corresponding to specific macroregions (e.g. cortex, thalamus, etc.) were independently shuffled over time.

### Experimental design and statistical analysis

Animal numbers were constrained by the labor intensity of the data collection methods yet comparable to previous publications using similar techniques and animal models ^15,54,96^. All control experiments were carried out in the same experimental cohort unless otherwise indicated in the main text. Sessions with degraded cranial windows or excessive virtual burrow motion were excluded from the analyses.

For all displayed maps, two-tailed one-sample t-tests were used in the second-level analysis to identify significant activation per voxel, taking individual mice as data points to avoid pseudoreplication. To account for the number of tests performed, a multiple comparison correction was implemented based on the Benjamini-Krieger-Yekutieli two-stage procedure to control the false discovery rate (FDR), setting a FDR < 0.15.

Unless otherwise indicated, group results for time series are displayed as mean and standard error across mice, and all box and whisker plots represent the median, interquartile range (box), and 5th to 95th percentile (whiskers) of the underlying distribution. The standard error of the mean corresponds to the standard deviation divided by the square root of the number of mice.

## Acknowledgements

We thank the lab members of the Mace lab, as well as S. Wiegert and N. Gogolla, for helpful discussions and feedback on the manuscript. Additional thanks go to N. Gogolla for providing us with viruses and resources for starting this project. We thank M. Reimers, P. Wallisch, M.X. Cohen and S. Solla for useful input regarding data analysis. We thank M. Mathis and S. Schneider for early access to the CEBRA code and support for implementing it with fUS data. We thank G. Montaldo and A. Urban for technical support with fUS data acquisition. We thank J. Deussing for providing the NAT-Cre mice, and the MPI-BI and UMG animal facilities and caretakers for their continued support. We thank the MPI-BI Imaging Facility and Histolab for assistance with histology and imaging. We thank J. Kuhl for help with illustrations. J.M.M.P. received support from the Joachim Hertz Foundation. All authors were funded by the Max Planck Society, the Else Kroner Fresenius Foundation through the Else Kroner Fresenius Center for Optogenetic Therapies, the Niedersachsisches Ministerium für Wissenschaft und Kultur through zukunft.niedersachsen, and the Deutsche Forschungsgemeinschaft (DFG, German Research Foundation) through Germany’s Excellence Strategy EXC 2067/1-390729940, the Project number 537609931 (FOR 5807, project 1) and the Collaborative Research Center 1690 (Project B05).

## Author Contributions

J.M.M.P. and E.M. conceptualized the study. J.M.M.P., J.L.M., P.W and E.M. contributed to the methodology. J.M.M.P., J.L.M., P.W., B.R.A., A.A. and L.B. carried out the investigation and provided curated data. J.M.M.P. analyzed the data. A.A. trained and apply the CEBRA model. J.M.M.P. generated the visualizations. J.M.M.P. and E.M. wrote the original draft and all authors contributed to the final version. E.M. supervised the project and acquired the funding.

## Data and code availability

All data and code will be made available on public repositories upon publication.

