## Supplementary figures and Supp Table 1 for "Arousal elicits a brain-wide hemodynamic wave independent of locus coeruleus noradrenergic tone"

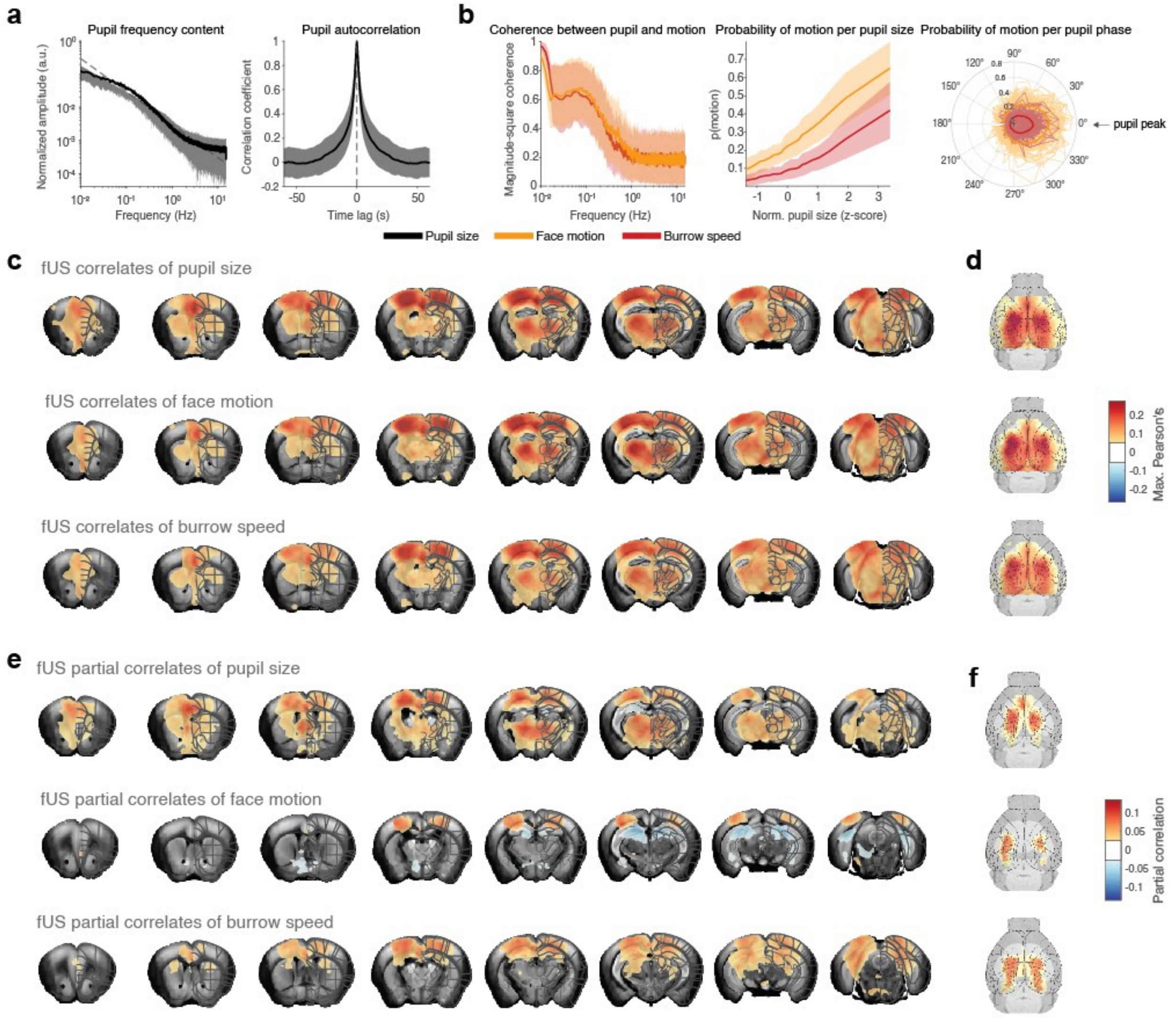


**Supp. Figure 1. Pupil and motion reflect the same underlying spontaneous arousal fluctuations, but pupil is more explanatory of brain-wide fUS signal.**

**a.** Left: pupil frequency content. Right: pupil autocorrelation. **b.** Left: coherence between pupil and burrow and face motion. Center: probability of motion as a function of normalized pupil size. Right: probability of motion as a function of pupil phase. **c.** Brain-wide maximum correlation map of pupil size (*top*), face motion (*middle*) and virtual burrow speed (*bottom*). **d.** Cortical views of the maps in **c**. **e.** As **c.**, but for brain-wide partial correlation maps. **f.** Cortical views of the maps in **e**.

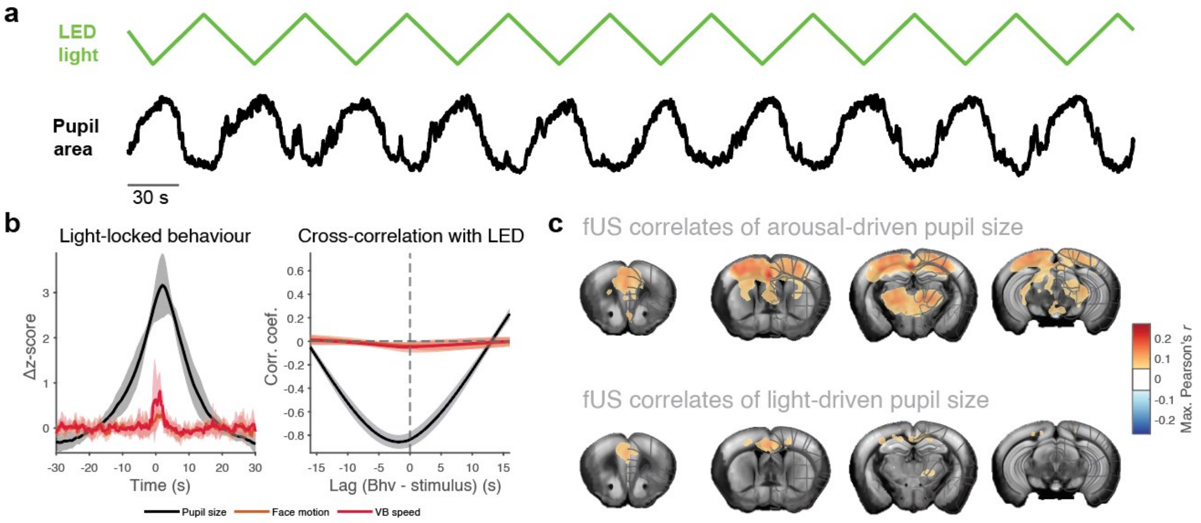


**Supp. Figure 2. Pupillary change alone does not evoke the arousal pattern.**

**a.** Ramping light stimulus used to trigger the photopupillary reflex (*top*) and representative associated pupil size trace (*bottom*). **b.** Average behavioral metrics locked to the timepoint of maximum illumination (*left*) and cross-correlation between the light stimulus and the behavioral traces (*right*). **c.** Brain-wide 0-lag correlation map of pupil size during constant illumination (*top*) and during ramping light stimulation (*bottom*) with fUS signal, with fine brain regions overlaid

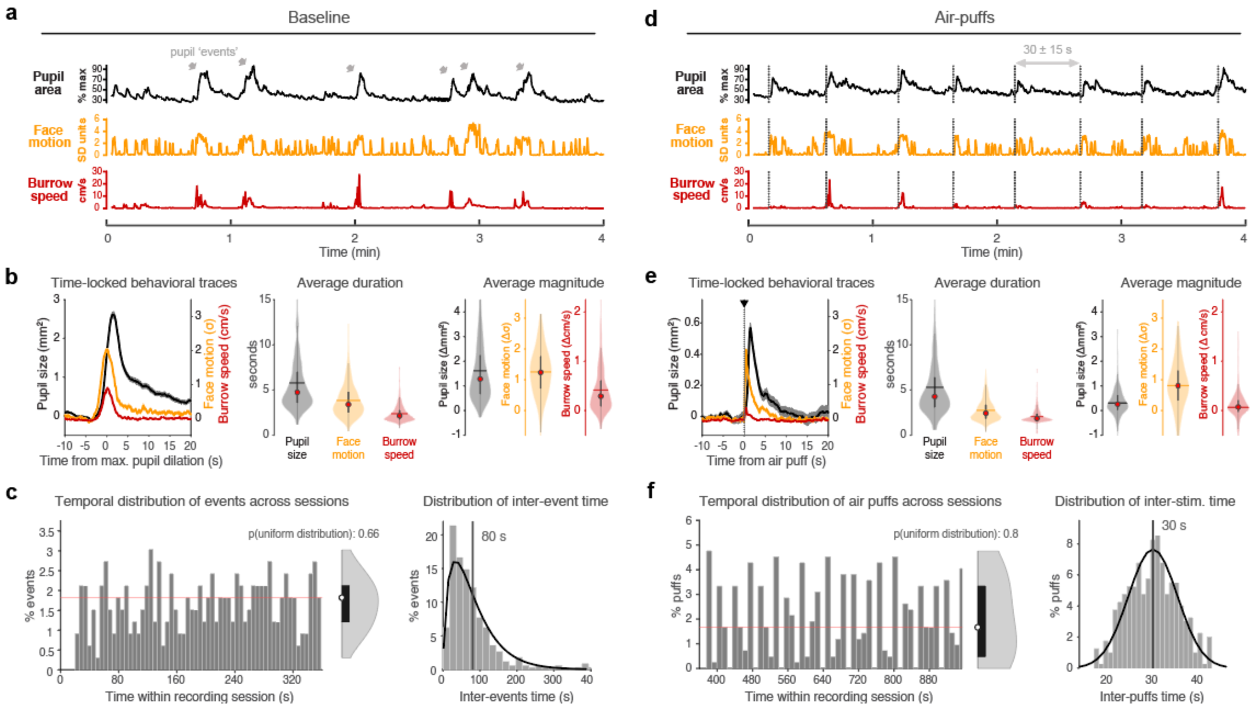


**Supp. Figure 3. Identification and behavioral characterization of spontaneous and evoked arousal transitions.** **a.** Example time series of behavioral metrics during a stimulus-free recording, with spontaneous arousal events highlighted. **b.** Left: average time-locked behavioral metrics during spontaneous arousal transitions. Center: distribution of each behavioral metric bout duration (full-width at half-maximum) across mice. Right: distribution of each behavioral metric peak value across mice. **c.** Left: probability of spontaneous pupil events throughout the typical recording session length. Center: probability distribution for the spontaneous events occurrence, including the probability of this distribution being uniform (calculated via Kolmogorov-Smirnov test). Right: histogram of the inter-event times, with average time highlighted. **d.** Example time series of behavioral metrics during an air puff recording, with the air puffs highlighted. **e.** As in **b**, but for stimulus-locked behavioral metrics during evoked arousal transitions. **f.** As in **c**, but for the probability of pseudorandomized air puffs throughout the typical recording session length (*left*, *center*) and for inter-stimulus times (*right*).

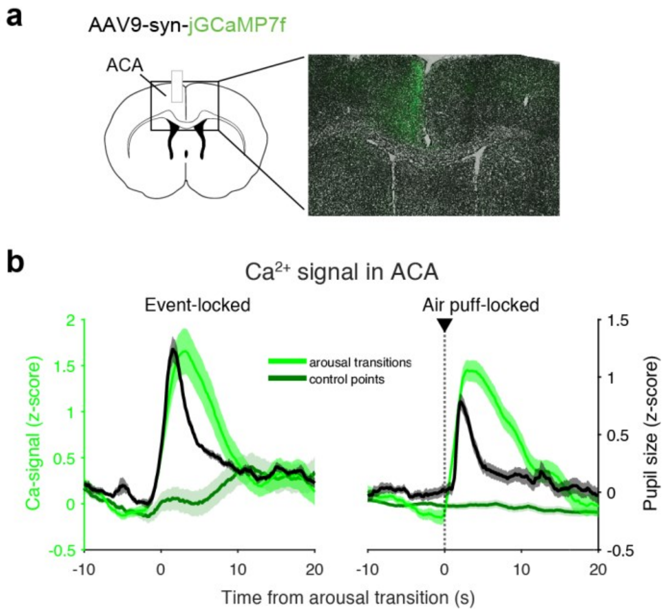


**Supp. Figure 4. Fiber photometry signal peaks upon spontaneous and evoked arousal.**

**a.** Diagram of the viral injection and fiber insertion for fiber photometry, and example confocal microscopy images showing GCaMP expressions in anterior cingulate cortex. **b.** Average $Ca^{2+}$ trace in anterior cingulate cortex upon spontaneous pupil events (*left*) and in response to air puffs (*right*), versus average $Ca^{2+}$ trace around control (random) timepoints. ACA: anterior cingulate area.

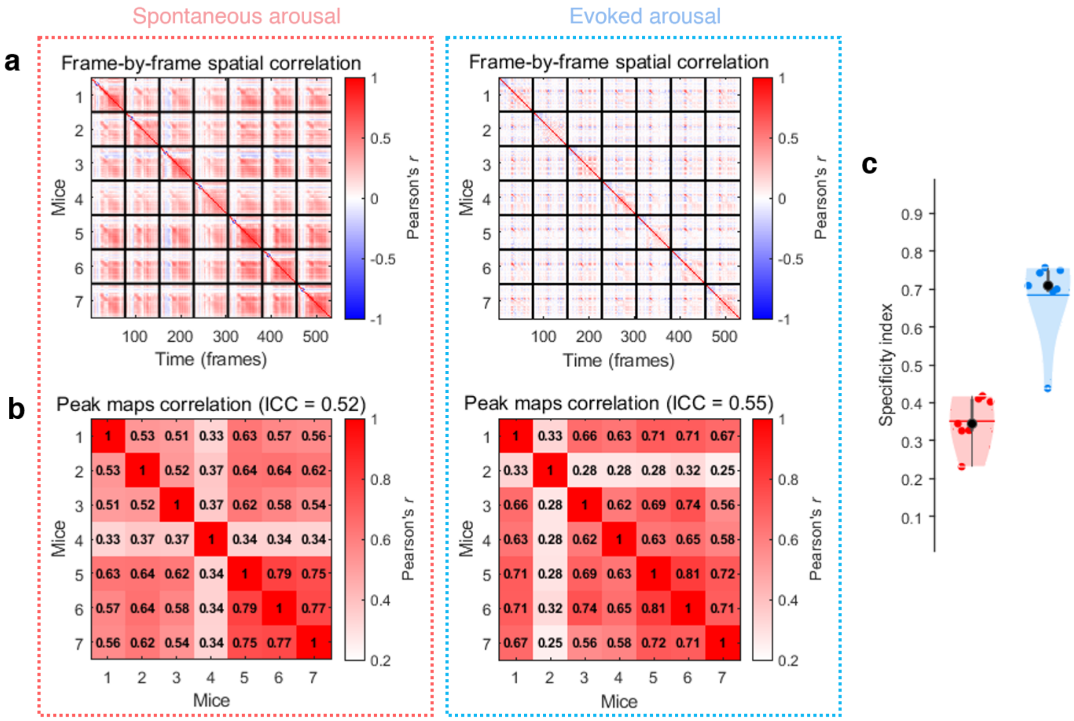


**Supp. Fig. 5. Replicability of the arousal responses across mice.**

**a.** Spatial correlation between each frame of the spontaneous (*left*) and evoked (*right*) arousal responses of each mouse. The checkboard pattern suggests that the peak response is consistent across mice. **b.** Spatial correlation between the peak amplitude maps for spontaneous (*left*) and evoked (*right*) arousal responses across mice. **c.** Specificity index of spontaneous and evoked arousal across mice, defined as (correlation with correct template) – (correlation with incorrect template), where the correct template is the average map of the arousal response and the incorrect template is the average map of the pre-arousal baseline. ICC: inter-class correlation.

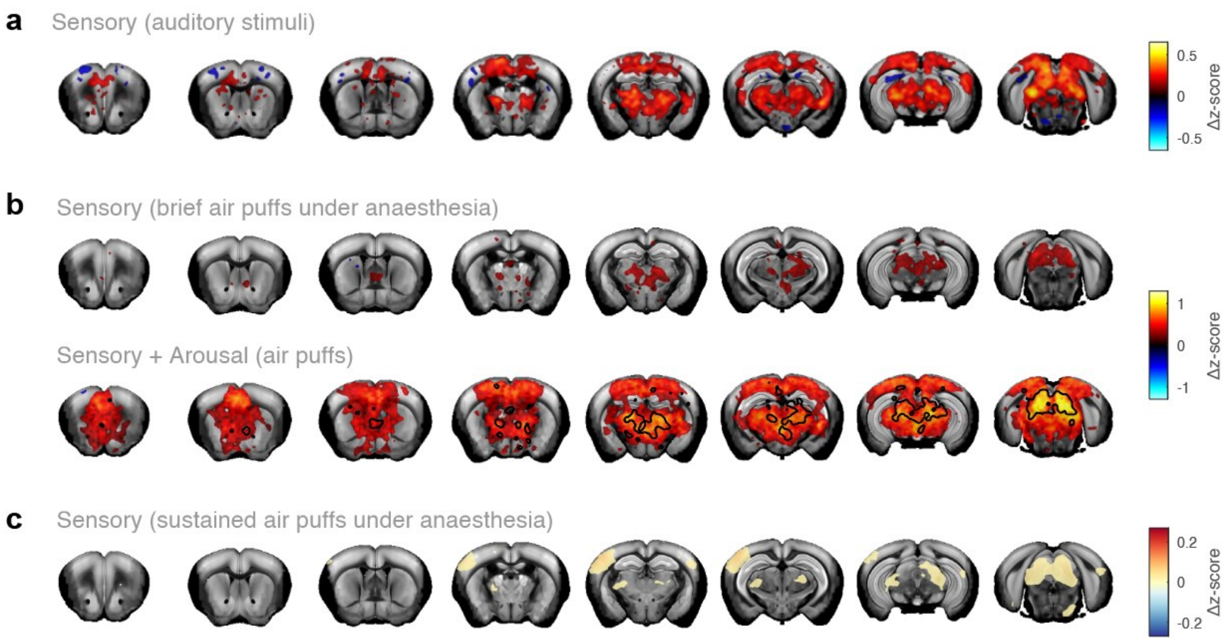


**Supp. Figure 6. Evoked arousal response is specific for awake stimulation.**

**a** Average brain-wide fUS response amplitude to white noise, an arousing auditory stimuli (399 stimuli, 7 mice).**b.** Top: average brain-wide fUS response amplitude for brief air puffs under anesthesia (560 puffs, 7 mice). Bottom: average brain-wide fUS response amplitude for brief air puffs during wakefulness (419 puffs, 7 mice), with the contour of regions still significant under anesthesia overlined. **c.** correlation map for sustained air puffs under anesthesia (28 sessions, 7 mice)**.**

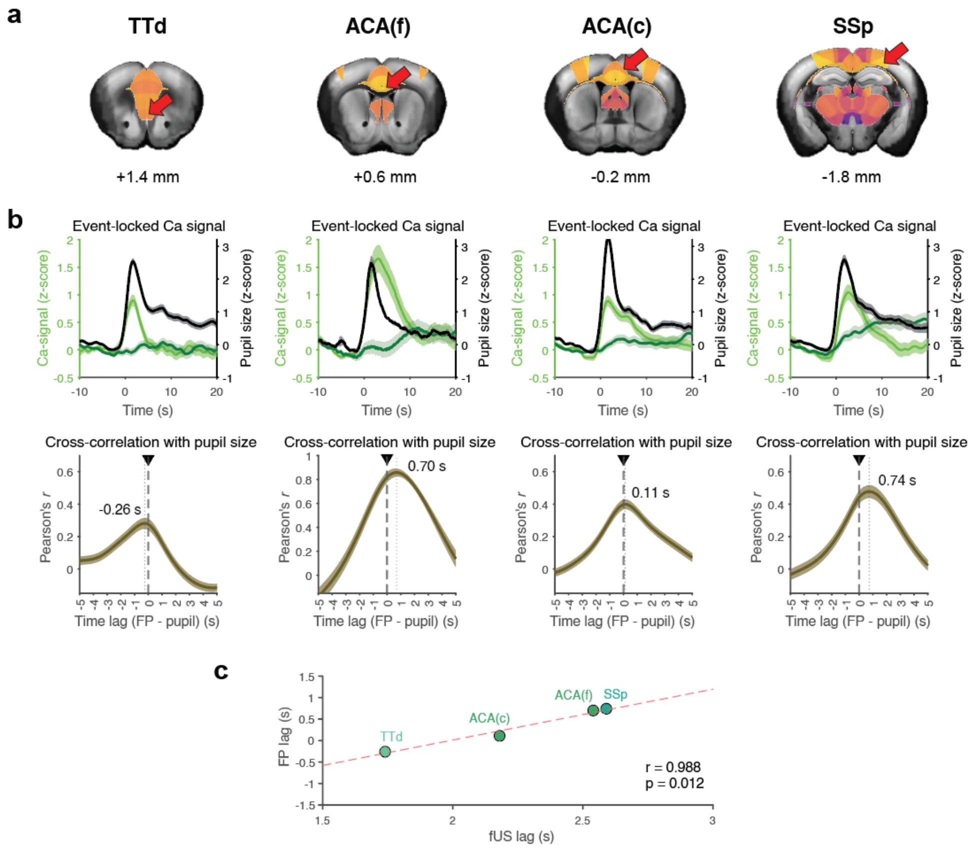


**Supp. Figure 7. Fiber photometry signal keeps the same temporal relationship to pupil size as fUS signal.**

**a.** Coronal sections showing the selected regions and their average order of activation as in **Fig. 3**. **b.** Top: average $Ca^{2+}$ traces upon spontaneous pupil events. Bottom: average cross-correlation between $Ca^{2+}$ traces and pupil traces during spontaneous events, used to calculate the delay between both traces. **c.** Scatterplot comparing the fUS and FP average temporal dynamics. TTd: dorsal taenia tecta; ACA: anterior cingulate area (f: frontal, c: caudal); SSP: primary somatosensory cortex.

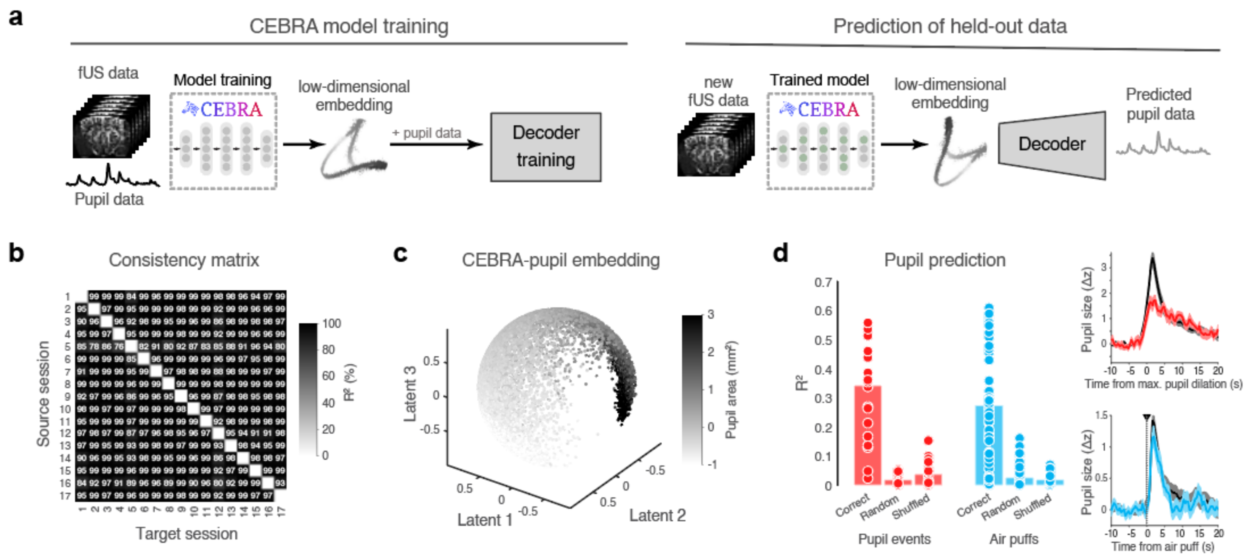


**Supp. Figure 8. CEBRA embedding trained on spontaneous fUS signal generalizes to evoked arousal.**

**a.** Schematic of the CEBRA approach. Parcellated fUS data and concurrent pupil size from concatenated baseline (stimulus-free) recordings (*n* = 14 sessions, *N* = 7 mice) were used to train CEBRA and generate low-dimensional representations of the data structure (latent embeddings), which in turn were used to train a kNN decoder. The trained model was used to project new recordings from both stimulus-free (*n* = 4 sessions, *N* = 4 mice) and air puffs (*n* = 21 sessions, *N* = 7 mice) conditions into the embedding, and the trained decoder was used to predict their associated pupil trace. **b.** Consistency matrix showing $R^2$ values after fitting a linear model between embeddings of independent session pairs. **c.** Latent embedding from CEBRA, plotted using 3 latent dimensions, where each dot represents one timepoint, color-coded by associated pupil size. **d.** Left: barplots representing the average $R^2$ of the pupil prediction generated by different CEBRA models: trained with correct pupil traces, trained with randomly-permuted pupil traces, and trained with shuffled-across-sessions pupil traces. Right: average pupil trace predicted from the "correct" model for both conditions.

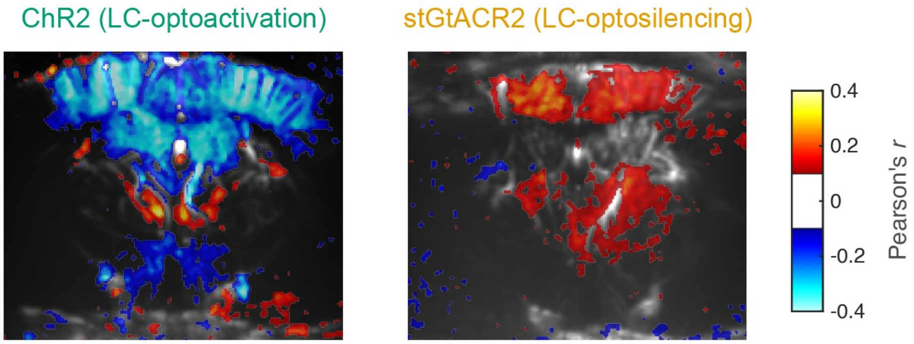


**Supp. Figure 9. LC optogenetic stimulation may induce spatially heterogeneous responses in caudal regions.** Example 2D-fUS coronal slices of optostimulation correlates at the caudal end of the imaging field-of-view for the optogenetic cohorts (thresholded for $r > 0.1$), suggesting differential effects of optostimulation in the thalamus.

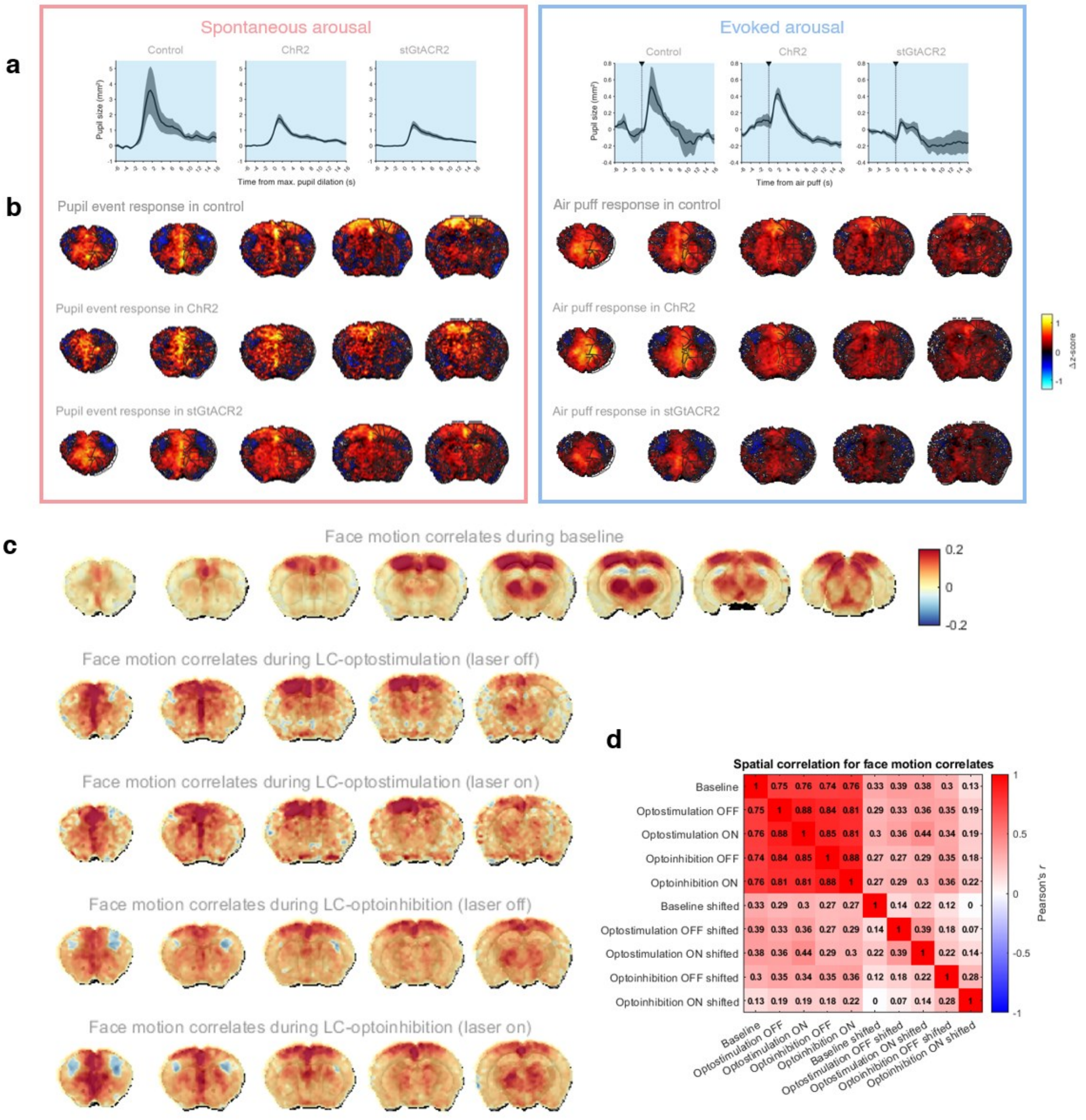


**Supp. Figure 10. Sustained opto-manipulations preserves arousal responses.**

**a.** Average time-locked pupil size during spontaneous (*left*) and evoked (*right*) arousal transitions. **b.** Normalized (z-scored) average brain-wide fUS response amplitude for spontaneous pupil events (*left*) and air puffs (*right*) during concurrent opto-manipulation for all three optomanipulation mice cohorts**. c.** Brain-wide maximum correlation maps of face motion with fUS signal for different mice cohorts. **d.** Matrix of spatial similarity (spatial Pearson's correlation coefficient) between the correlation maps for each mice cohort, including control maps where the face motion regressor has been shifted to eliminate the temporal alignment with fUS signal.

### Supplementary table 1: list of common arousal regions as per Allen Brain Atlas demarcation

| | | |
|---|---|---|
| Primary somatosensory area, lower limb | Primary somatosensory area, trunk | Primary somatosensory area, upper limb |
| Anterior cingulate area, dorsal part | Anterior cingulate area, ventral part | Anterior visual area |
| Anteromedial visual area | Primary visual area, anterior lateral | Primary visual area, anterior medial |
| Primary visual area, posterior medial | Posteromedial visual area | Rostrolateral visual area |
| Prelimbic area | Infralimbic area | Restrosplenial area, lateral agranular part |
| Retrosplenial area, dorsal part | Retrosplenial area, ventral part | Taenia tecta, dorsal part |
| Dorsal peduncular area | CA2 subfield, dorsal part | Induseum griseum |
| Postsubiculum | Area prostriata | Lateral septal nucleus, rostral (rostroventral) part |
| Septofimbrial nucleus | Triangular nucleus of septum | Anterodorsal nucleus |
| Anteromedial nucleus, dorsal part | Anteromedial nucleus, ventral part | Anteroventral nucleus of thalamus |
| Interanterodorsal nucleus of the thalamus | Intergeniculate leaflet of the lateral geniculate complex | Ventral part of the lateral geniculate complex |
| Intermediate geniculate nucleus | Subgeniculate nucleus | Central lateral nucleus of the thalamus |
| Central medial nucleus of the thalamus | Paracentral nucleus | Posterior intralaminar thalamic nucleus |
| Rhomboid nucleus | Ethmoid nucleus of the thalamus | Posterior complex of the thalamus |
| Lateral posterior nucleus of the thalamus | Suprageniculate nucleus | Posterior limiting nucleus of the thalamus |
| Intermediodorsal nucleus of the thalamus | Mediodorsal nucleus of thalamus | Perireunensis nucleus |
| Submedial nucleus of the thalamus | Parataenial nucleus | Dorsal part of the lateral geniculate complex, core |
| Dorsal part of the lateral geniculate complex, ipsilateral zone | Medial geniculate complex, medial part | Medial geniculate complex, ventral part |
| Peripeduncular nucleus | Subparafascicular nucleus, parvicellular part | Posterior triangular thalamic nucleus |
| Ventral anterior-lateral complex of the thalamus | Ventral medial nucleus of the thalamus | Ventral posterolateral nucleus of the thalamus |
| Ventral posteromedial nucleus of the thalamus | Lateral habenula | Medial habenula |
| Zona incerta | Parasubthalamic nucleus | Medial mammillary nucleus, lateral part |
| Medial mammillary nucleus, median part | Supramammillary nucleus | Posterior hypothalamic nucleus |
| Central linear nucleus raphe | Interfascicular nucleus raphe | Interpeduncular nucleus, rostral |
| Interpeduncular nucleus, rostrolateral | Rostral linear nucleus raphe | Pedunculopontine nucleus |
| Substantia nigra, compact part | Anterior tegmental nucleus | Dorsal terminal nucleus of the accessory optic tract |
| Edinger-Westphal nucleus | Medial accesory oculomotor nucleus | Medial terminal nucleus of the accessory optic tract |
| Midbrain reticular nucleus, anterior lateral | Midbrain reticular nucleus, anterior medial | Midbrain reticular nucleus, posterior lateral |
| Midbrain reticular nucleus, posterior medial | Paranigral nucleus | Periaqueductal gray, anterior part |
| Periaqueductal gray, medial dorsal | Periaqueductal gray, medial lateral | Periaqueductal gray, medial ventral |

| | | |
|---|---|---|
| Periaqueductal gray, posterior dorsal | Periaqueductal gray, posterior lateral | Precommissural nucleus |
| Nucleus of Darkschewitsch | Interstitial nucleus of Cajal | Supraoculomotor periaqueductal gray |
| Anterior pretectal nucleus | Nucleus of the posterior commissure | Retroparafascicular nucleus |
| Red nucleus | Substantia nigra, reticular part | Superior colliculus, deep layers, posterior |
| Superior colliculus, intermediate layers, anterior lateral | Superior colliculus, intermediate layers, anterior medial | Superior colliculus, intermediate layers, posterior lateral |
| Superior colliculus, superficial layers, anterior lateral | Superior colliculus, superficial layers, anterior medial | Superior colliculus, superficial layers, posterior lateral |
| Ventral tegmental area | Inferior colliculus, external nucleus | Nucleus sagulum |
| Parabigeminal nucleus | Pontine reticular nucleus | |
